# The SensorOverlord predicts the accuracy of measurements with ratiometric biosensors

**DOI:** 10.1101/2020.01.31.928895

**Authors:** Julian A. Stanley, Sean B. Johnsen, Javier Apfeld

**Affiliations:** Biology Department, Northeastern University, Boston, MA, 02115, USA

## Abstract

Two-state ratiometric biosensors change conformation and spectral properties in response to specific biochemical inputs. Much effort over the past two decades has been devoted to engineering biosensors specific for ions, nucleotides, amino acids, and biochemical potentials. The utility of these biosensors is diminished by empirical errors in fluorescence-ratio signal measurement, which reduce the range of input values biosensors can measure accurately. Here, we present a formal framework and a web-based tool, the SensorOverlord, that predicts the input range of two-state ratiometric biosensors given the experimental error in measuring their signal. We demonstrate the utility of this tool by predicting the range of values that can be measured accurately by biosensors that detect pH, NAD^+^, NADH, NADPH, histidine, and glutathione redox potential. The SensorOverlord enables users to compare the predicted accuracy of biochemical measurements made with different biosensors, and subsequently select biosensors that are best suited for their experimental needs.

## Introduction

Genetically encoded two-state ratiometric biosensors have revolutionized our ability to monitor a wide variety of biochemical species^1–8^. The development of these biosensors has enabled the visualization in real-time of the biochemical properties of live animals using fluorescence-ratio microscopy. However, the potential of these biosensors has not been fully realized because the empirical imprecision of their fluorescence-ratio signal measurements reduces the range of biochemical input values those biosensors can measure accurately.

The capacity to make accurate measurements with sensors is important because it enables observers to make confident predictions about the state of a system. Using a thermometer that makes inaccurate temperature measurements can lead to incorrect predictions about the state of a physical system; for example, in predicting whether water will be a solid, a liquid, or a gas. Similarly, using a genetically encoded biosensor that makes inaccurate measurements can lead to incorrect predictions about the state of a biological system.

In our previous work in the nematode *C. elegans*, we deployed a mathematical framework that enabled us to map the fluorescence-ratio signal of the roGFP1-R12 biosensor into glutathione redox potential (*E*_*GSH*_) values using prior information about our microscope’s properties and the biosensor’s spectral and biochemical properties^9,10^. Here, we extend that framework to determine how the precision of our fluorescence-ratio signal measurements with the roGFP1-R12 biosensor constrains the range of *E*_*GSH*_ values that can be measured accurately. We then generalize this extended framework for all two-state ratiometric biosensors with known spectral and biochemical properties. We demonstrate the utility of this new framework by: (i) determining the range of *E*_*GSH*_ values that we can measure accurately in live *C. elegans* with the roGFP1-R12 biosensor; (ii) quantifying how much that range of *E*_*GSH*_ values is expanded by increasing the precision of our imaging and image-analysis methods; (iii) identifying which biosensors are best suited for measuring accurately different ranges of *E*_*GSH*_, pH, and the concentrations of nucleotides and amino acids; (iv) identifying underused biosensors; and (v) identifying where new biosensors are needed.

To help the community identify biosensors that are well-suited for their experimental needs, we developed a web-based tool, the SensorOverlord (https://www.sensoroverlord.org), that implements all of these analyses with a user-friendly interface.

## Results

### Predicting the accuracy of a glutathione redox potential biosensor

The human and *C. elegans* proteomes contain ~210,000 cysteine residues^11,12^ and ~15% of these cysteines are reversibly oxidized^13^. These protein networks can be understood as markets where cysteines in proteins buy (*reduction*) and sell (*oxidation*) pairs of electrons only via a central broker, the abundant glutathione tripeptide^9^, resulting in a single price for trading electrons that determines the oxidation of all cysteines in the network (Figure 1a). In chemical terms, this price is the glutathione redox potential (*E*_*GSH*_): the Nernst potential that quantifies the balance between reduced and oxidized glutathione species. Measuring *E*_*GSH*_ is critical because cysteine oxidation modulates the function of hundreds of cytosolic proteins^14–20^ which regulate a wide variety of cellular processes^20,21^. The mechanisms that regulate *E_GSH_ in vivo* remained largely unexplored until the development of the *E*_*GSH*_-specific roGFP-family of genetically-encoded biosensors^10,22,23^. These GFP-derived biosensors include two cysteines that form a (reversible) intramolecular disulfide bond upon oxidation, resulting in spectral changes that can be quantified via fluorescence-ratio microscopy (Figure 1b)^8^. We previously used the roGFP1-R12 biosensor to measure *E*_*GSH*_ in live *C. elegans*^9^. To map fluorescence-ratio (*R*) measurements into *E*_*GSH*_ values, we determined three conversion factors that quantify the properties of our imaging microscope and the spectral differences between the reduced and oxidized states of the biosensor (Supplementary Note 1). Measuring *E*_*GSH*_ instead of *R* enabled us to make predictions about how the oxidation state of the network of cysteines trading electrons with glutathione is influenced by genetic determinants and environmental factors^9^. However, those predictions require that *E*_*GSH*_ be measured accurately. Therefore, we set out to determine how the precision of our fluorescence-ratio microscopy influenced the range of *E*_*GSH*_ values we could measure accurately.

**Figure 1.**
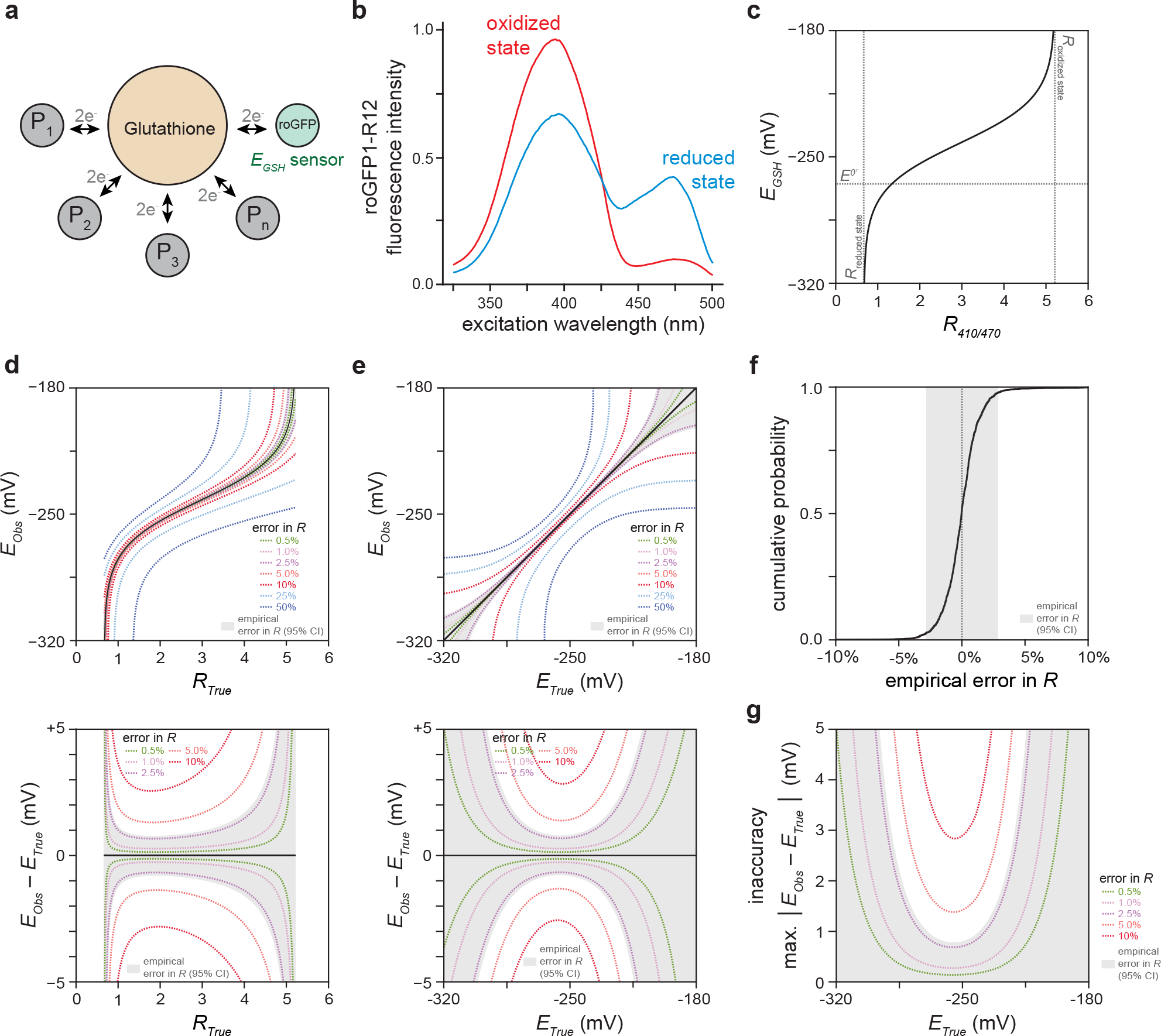
Determining the range of glutathione redox potential *E*_*GSH*_ values we can measure accurately with the roGFP1-R12 biosensor. **a.** Glutathione redox potential (*E*_*GSH*_) directs the oxidation of cysteines in hundreds of proteins in the same direction, resulting in their concerted regulation. **b.** The reduced and oxidized states of the roGFP1-R12 biosensor have different fluorescence spectra^8^, enabling *E* measurement via fluorescence-ratio (*R*) microscopy. **c.** The conversion map from *R* to *E*_*GSH*_ is highly nonlinear. *R*_*reduced state*_ and *R*_*oxidized state*_ refer to the ratiometric emission of ensembles of reduced and oxidized biosensors, respectively. *E*^*0’*^ is the standard midpoint potential of the biosensor. **d.** The top panel shows how measurement errors in *R* cause observed *E*_*GSH*_ values (*E*_*Obs*_) to differ from the true *E*_*GSH*_ values (*E*_*True*_) that would be observed if *R* was measured with no error (*R*_*True*_). The bottom panel shows how the size of an *E*_*GSH*_ error (*E*_*Obs*_ – *E*_*True*_) depends not only on the size of the error in *R* but also on the value of *R*. Each dotted curve corresponds to a different fold-change error in *R*. The shaded region corresponds the interval encompassing 95% of the predicted *E*_*Obs*_ values for each *R*_*True*_ value, given our empirical error in *R.* **e.** Transforming the map from *R*_*True*_ to *E*_*True*_ in the top and bottom panels shown in **d** produces plots showing how errors in *R* influence the map from *E*_*True*_ to *E*_*Obs*_ (top panel) and how the size of an *E*_*GSH*_ error depends not only on the size of the error in *R* but also on the value of *E*_*True*_ (bottom panel). Each dotted curve corresponds to a different fold-change error in *R*. The shaded region shows the interval encompassing 95% of the predicted *E*_*Obs*_ values for each *E*_*True*_ value, given our empirical error in *R.* **f.** Cumulative distribution of the empirical fold error in *R* in live *C. elegans* expressing the roG-FP1-R12 biosensor in the cytosol of the anterior (pm3) muscles of the pharynx, the feeding organ. This error distribution was obtained by aggregating with equal weight the empirical fold error in *R* of five separate experiments (see Supplementary Note 3). 95% of the errors in *R* fall within the interval (−2.8%, +2.8%), shown shaded in gray. This interval quantifies the precision of our fluorescence-ratio measurements. **g.** *E*_*GSH*_ measurement inaccuracy (the maximum absolute difference between *E*_*True*_ and *E*_*Obs*_) decreases with increased precision of *R* measurement. Each dotted curve corresponds to a different precision of *R* measurement. The shaded region shows the interval encompassing 95% of the predicted *E*_*GSH*_ measurement inaccuracies for each *E*_*True*_ value, given our empirical error in *R.*

We first modeled how errors in fluorescence-ratio measurement influenced *E*_*GSH*_ errors. The conversion map from *R* to *E*_*GSH*_ is highly nonlinear (Figure 1c). As a result, the size of an *E*_*GSH*_ error depends not only on the size of the error in *R* but also on the value of *R* (Figure 1d): as *R* approaches its lower and upper bounds *E*_*GSH*_ errors increase rapidly (Supplementary Note 2). Thus, even a small difference between observed and true *R* values (*R*_*Obs*_ and *R*_*True*_, respectively) can lead to a large difference between observed and true *E*_*GSH*_ values (*E*_*Obs*_ and *E*_*True*_, respectively) (Figure 1d).

We then determined the size of our fluorescence-ratio measurement errors. We quantified the precision of our fluorescence-ratio measurements in live *C. elegans* expressing the roGFP1-R12 biosensor in the cytosol of the muscles of the pharynx, the feeding organ. This retrospective analysis of 10,572 images showed that our errors in *R* were proportional to *R*—that is, *R*_*Obs*_ = *R*_*True*_ * (1 + *error*) (Supplementary Note 3). Within a given experiment, the size of the relative error in *R* was invariant over the range of all possible *R* values (Supplementary Note 3). The size of the relative error in *R*, however, varied up to three-fold between experiments (Supplementary Note 3). Differences in the proportion of animals moving during imaging accounted for most of the variation in the relative error in *R* across experiments (S.B.J., J.A.S., and J.A., manuscript in preparation). Our analysis indicated that, in a typical experiment, the median relative error in *R* was zero and 95% of the relative errors in *R* were in the interval (−2.8%, +2.8%) (Figure 1f). These 95% confidence bounds quantified the precision of our fluorescence-ratio measurements.

Last, we determined how the empirical precision of our fluorescence-ratio measurements influenced the accuracy of individual *E*_*GSH*_ observations. Knowing the precision of our *R* measurements enabled us to determine the 95% confidence bounds of *E*_*Obs*_ as a function of *R*_*True*_ (Figure 1d). Converting *R*_*True*_ into *E*_*True*_ produced a map of how the 95% confidence bounds of *E*_*Obs*_ varied as a function of *E*_*True*_ (Figure 1e). The maximum absolute difference between *E*_*True*_ and either the upper or lower 95% confidence bound of *E*_*Obs*_ represents the inaccuracy of our *E*_*GSH*_ measurements (Figure 1g). Our mathematical modeling indicated that the precision of *R* measurements, the biochemical and biophysical properties of the biosensor, and the choice of excitation wavelengths used in our experiments all influenced the *E*_*GSH*_ values that we could measure most accurately (Supplementary Note 4). *E*_*GSH*_ inaccuracy rapidly increased as *E*_*True*_ moved farther away from that value.

This analysis enabled us to extract the range of *E*_*GSH*_ values that our biosensor was well-suited to measure at a given level of *E*_*GSH*_ inaccuracy (Figure 1g). For example, the range of *E*_*Obs*_ values we could measure with an inaccuracy of 2 mV was between −284 and −234 mV. This range encompassed all *E*_*GSH*_ values we observed in wild-type nematodes under normal conditions (−278 to −262 mV) and under oxidative stress (−278 to −250 mV)^9^, indicating that our experimental set up was well-suited to measure the *E*_*GSH*_ values that *C. elegans* feeding muscles exhibited *in vivo*: 95% of the individual *E*_*GSH*_ observations deviated from their true value by less than 2 mV.

### Balancing the need for accurate measurements with the constraints of microscopy

Our analytical framework provides a criterion for determining if it is possible to measure *E*_*GSH*_ accurately. Scientific needs demand accurate observations, but experimental approaches constrain the extent to which observations can be made accurately. The trade-off between these scientific and experimental constraints can be visualized in a phase diagram (Figure 2). The precision of *R* measurements determines the range of *E*_*GSH*_ values that is possible to measure at a specific inaccuracy level (Figure 2). For values outside that range, it is impossible to guarantee that an observation will be accurate. Scientific needs impose a maximum tolerable inaccuracy beyond which observations are too inaccurate and, therefore, not useful. Together, these constraints determine whether it is possible to measure *E*_*GSH*_ accurately (Figure 2).

**Figure 2.**
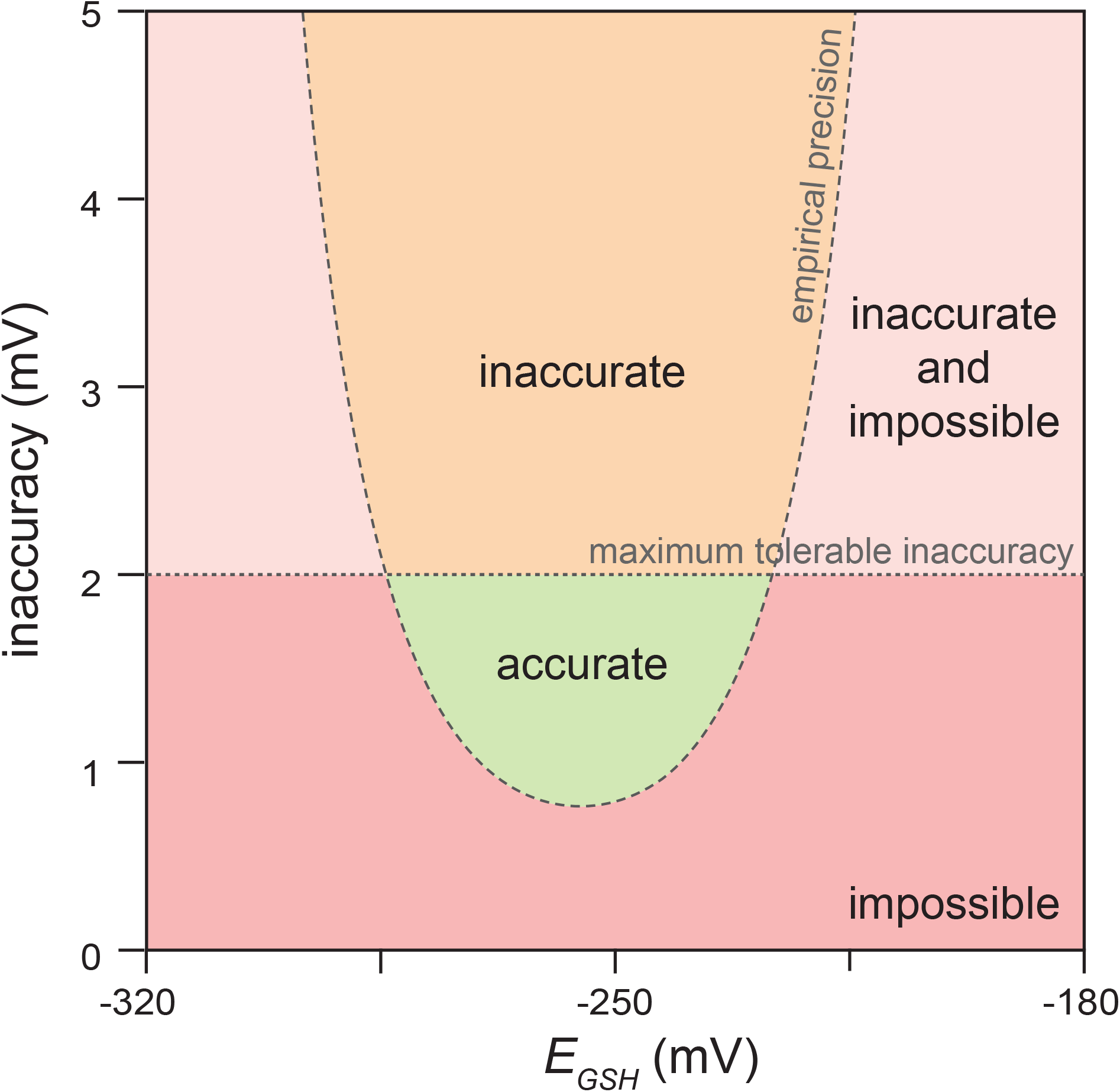
Balancing the need for accurate measurements with the constraints of microscopy. The empirical precision of our *R* measurements determines the range of *E*_*GSH*_ values that is possible to measure at a specific inaccuracy level. Values outside that range are impossible to measure accurately (red and light red regions). Scientific needs impose a maximum tolerable inaccuracy beyond which observations are too inaccurate and, therefore, not useful (light red and orange regions). Together, these constraints determine whether it is possible to accurately measure *E*_*GSH*_ (green region).

### Retrospectively increasing measurement accuracy with improved image analysis

To increase the range of *E*_*GSH*_ values that we could measure accurately, we set out to improve our image-analysis methods. Movement of live *C. elegans* during image acquisition lowers the precision of fluorescence-ratio measurements in individual pharyngeal muscles. In a typical experiment 21% of animals moved during imaging. We developed a new image-feature registration algorithm that corrects for displacement and deformation of the muscles along the anterior-posterior axis of the pharynx (S.B.J. and J.A., manuscript in preparation). This new image-analysis algorithm reduced the relative error in *R* along most positions in the pharynx, especially in the boundaries between adjacent muscles and in the muscles of the anterior and posterior bulbs. For example, in the pm7 muscles of the posterior bulb, the new algorithm reduced the interval with 95% of the relative errors in *R* from ±4.3% to ±2.6% in moving animals and from ±2.0% to ±1.9% in stationary animals. As a result, the new algorithm increased the accuracy with which we could measure *E*_*GSH*_ and thereby expanded the range of *E*_*GSH*_ values that we could measure accurately in past experiments (Figure 3a).

**Figure 3.**
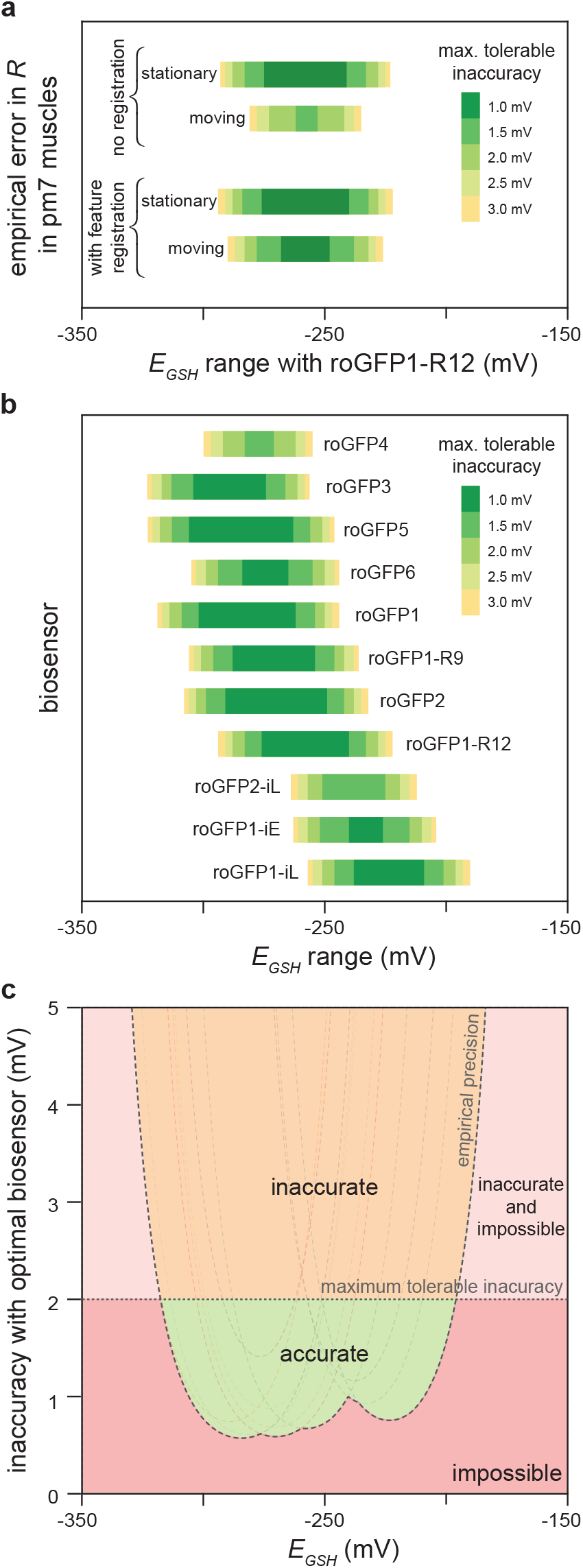
Predicted accuracy of glutathione redox potential biosensors. **a.** Predicted accuracy gains from improved image analysis in the pm7 (posterior) feeding muscles of live *C. elegans* expressing the roGFP1-R12 biosensor. Animals that moved during image acquisition showed a higher *R* measurement error than stationary animals. A feature-registration algorithm increased the precision of *R* measurements, retrospectively expanding the range of *E*_*GSH*_ values that we could measure accurately. The colored bars denote the range of *E*_*GSH*_ values where we have 95% confidence that an individual *E*_*GSH*_ observation would deviate from its true value by less than the error denoted by the color of the bar. **b.** Predictions of the ranges of *E*_*GSH*_ values that we expect to measure accurately in pm3 pharyngeal muscles with eleven roGFP-based biosensors given the empirical precision of our *R* measurements. Coloring of bars as in panel **a**. **c.** The empirical precision of our *R* measurements determines the range of *E*_*GSH*_ values that would be possible to measure at a specific inaccuracy level if we measured *E*_*GSH*_ in the pharyngeal muscles of live *C. elegans* with the most accurate roGFP biosensor for each *E*_*GSH*_ value. Values outside that range are impossible to measure accurately (red and light red regions). Scientific needs impose a maximum tolerable inaccuracy beyond which observations are too inaccurate and, therefore, not useful (light red and orange regions). Together, these constraints determine whether it is possible to accurately measure *E*_*GSH*_ with the eleven roGFP biosensors (green region). The dotted curves correspond to the predicted *E*_*GSH*_ inaccuracies of each of the eleven roGFP biosensors shown in **b**, given the precision of our *R* measurements.

### Comparing glutathione redox potential biosensors

We determined the ranges of *E*_*GSH*_ values that we could have measured accurately had we used different biosensors. Theoretical modeling indicated that the accuracy of a biosensor is influenced by the choice of wavelengths used for biosensor excitation, and by the biosensor’s dynamic range and midpoint-potential (*E*^*0’*^, the price point where a biosensor is 50% likely to sell its electrons) (Supplementary Note 4). These biosensor physical and chemical properties vary among all existing roGFP-based biosensors (Supplementary Note 5). We estimated the conversion factors that map fluores-cence-ratio measurements into *E*_*GSH*_ values for the eleven roGFP-based biosensors with known midpoint potentials and fluorescence spectra (Supplementary Note 5). This enabled us to determine the *E*_*GSH*_ inaccuracy we would expect to observe had we measured *E*_*GSH*_ in the feeding muscles of live *C. elegans* with each of those biosensors instead of roGFP-R12 (Figure 3b and Supplementary Note 5). This analysis enabled us to identify which biosensors would measure *E*_*GSH*_ most accurately under our experimental conditions: roGFP5 for *E*_*GSH*_ values below −297 mV, roGFP2 for *E*_*GSH*_ values from −296 mV to −258 mV, roGFP1-R12 for *E*_*GSH*_ values from −257 to mV to −240 mV, and roGFP1-iE for *E*_*GSH*_ values above −239 mV. We note that often many biosensors were predicted to have comparable accuracies (Figure 3b).

This analysis helped us identify underused biosensors. Neither roGFP3 nor roGFP5 has ever been used *in vivo*, yet we predict that these biosensors would be the most accurate biosensors for low *E*_*GSH*_ values such as those expected for the mitochondrial matrix. We currently disfavor roGFP5, even though this biosensor was predicted to be more accurate than roGFP3, because roGFP5 can potentially form more than one type of internal disulfide bridge due to its two additional cysteines; a better understanding of roGFP5’s biochemistry is warranted given its potential utility.

Comparison of the predicted accuracy of biosensors originally designed for similar purposes enabled us to identify the variables that explain why one biosensor was predicted to be more accurate than another (Supplementary Note 6). For example, both roGFP1-iE and roG-FP2-iL were designed to have higher midpoint potentials than previous roGFPs, making them more suitable for measuring the higher *E*_*GSH*_ values common in the endoplasmic reticulum. However, while roGFP1-iE has a higher midpoint potential than roGFP2-iL, it is predicted to be more inaccurate than roGFP2-iL even for measuring higher *E*_*GSH*_ values. The higher dynamic range of roGFP2-iL makes it a more accurate *E*_*GSH*_ biosensor than roGFP1-iE.

### Identifying where new glutathione redox potential biosensors are needed

We predicted the *E*_*GSH*_ inaccuracy that we would observe if we measured *E*_*GSH*_ in the feeding muscles of live *C. elegans* with the most accurate biosensor for each *E*_*GSH*_ value. Using a phase diagram, we visualized the trade-off between our scientific need for accuracy and the experimental constraints imposed by the precision of our *R* measurements and the properties of existing biosensors (Figure 3c). This analysis indicated that we lack biosensors well-suited to measure *E*_*GSH*_ values above −177 mV or below −337 mV with at least 10 mV accuracy.

### A general framework to predict the accuracy of two-state ratiometric biosensors

To establish a general criterion for determining whether a two-state biosensor is well-suited to measure its input accurately, we generalized the analysis framework for glutathione redox potential biosensors to all ratiometric two-state single-ligand-binding biosensors (Supplementary Notes 1, 5, 7). To demonstrate the utility of the generalized framework, we applied it to biosensors that measure pH and small molecules, including histidine, NAD^+^, NADH, and NADPH. For each biosensor with known affinity constant and fluorescence spectra, we derived the conversion factors that map its fluorescence-ratio to pH or ligand concentration (Supplementary Notes 8, 9). We then determined the pH and ligand concentration ranges that each biosensor would be well-suited to measure accurately given the precision of our *R* measurements and selecting optimal excitation or emission filters for each biosensor (Figures 4a-b and Supplementary Notes 8, 9).

**Figure 4.**
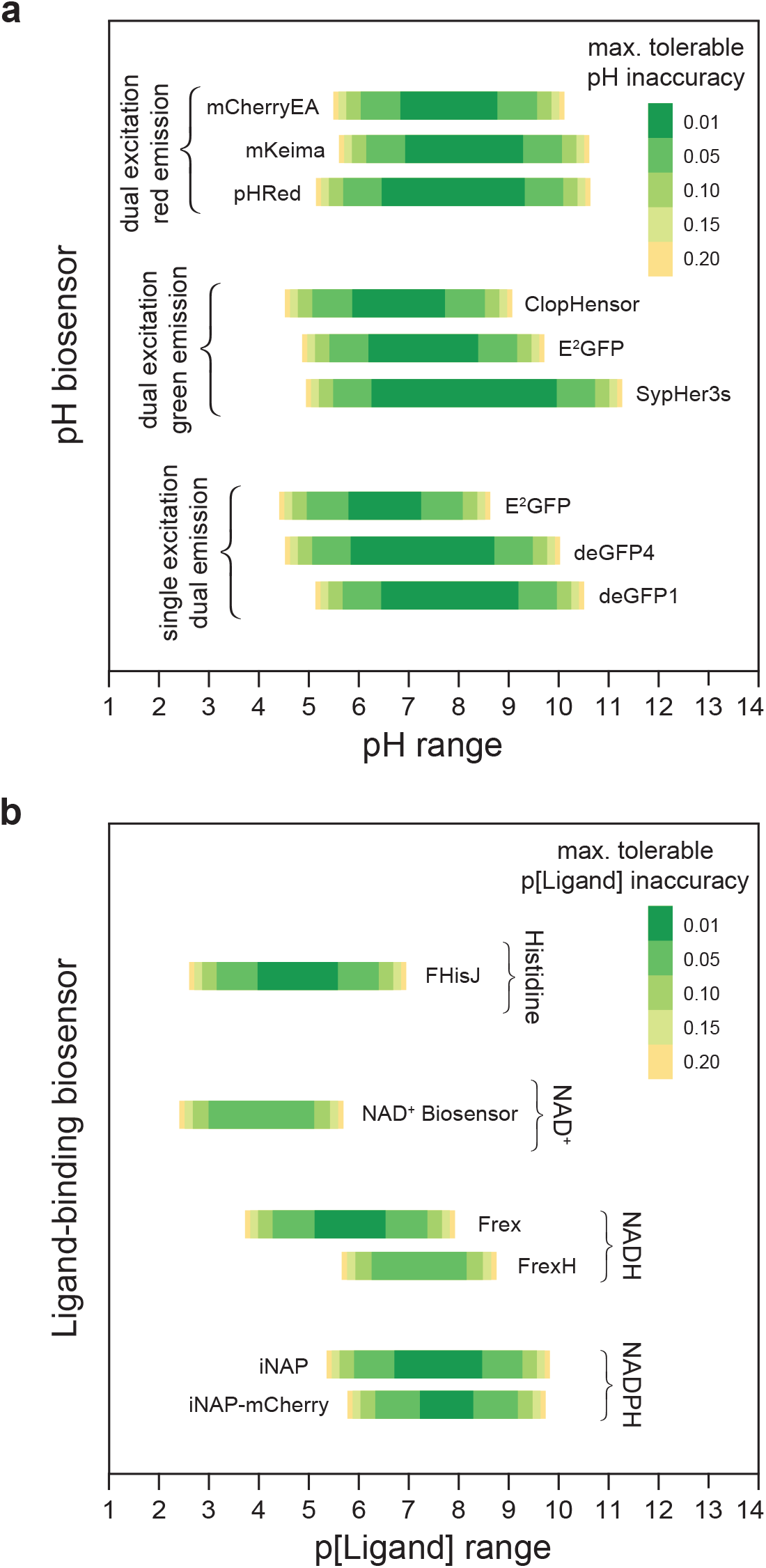
Predicted accuracy of pH and ligand-binding biosensors. Predictions of the ranges of pH (panel **a**), and histidine, NAD^+^, NADH, and NADPH values (panel **b**) that we expect to measure accurately in pm3 pharyngeal muscles with existing biosensors given the empirical precision of our *R* measurements and selecting optimal excitation or emission filters for each biosensor. The E^2^GFP biosensor can be used in two different modalities, dual-excitation green-fluorescence and single-excitation dual-emission. Differences in the predicted pH inaccuracy of this biosensor under each imaging modality arise from the differences between the values in each imaging modality of this biosensor’s overall dynamic range and dynamic range in the second wavelength (Supplementary Note 8). The colored bars denote the range of values of the biosensor’s biochemical input where we have 95% confidence that an individual observation would deviate from its true value by less than the error denoted by the color of the bar. p[Ligand] is the negative base 10 logarithm of the Molar concentration of the biosensor’s ligand.

Comparison of the predicted accuracy of nine ratiometric pH biosensors identified optimal biosensors for pH measurement with dual-excitation red-fluorescent pH biosensors, dual-excitation green-fluorescent pH biosensors, and single-excitation dual-emission pH biosensors (Figure 4a). The NADH-specific Frex biosensor^6^ had a higher predicted accuracy than the FrexH biosensor^6^, as a result of its higher dynamic range (Figure 4b). The NADPH-specific iNAP1 biosensor^7^ was predicted to more accurately measure NADPH concentration than the iNAP1-mCherry biosensor (Figure 4b). The iNAP1-mCherry biosensor sacrifices the iNAP1 dynamic range in one excitation band with pH-sensitive fluorescence, enabling pH-resistant NADPH measurement but lowering this biosensor’s accuracy.

### A web-based tool that predicts biosensor accuracy

To help the community find biosensors that are well-suited for their experimental needs, we developed the SensorOverlord toolkit. This open-source S4 class-based R package implements all the analyses described here. We also built a user-friendly web application, available at http://www.sensoroverlord.org (Figure 5). The SensorOverlord toolkit enables users to model how the precision of their fluores-cence-ratio signal measurements and their microscopy configuration constrain the range of input values that their biosensor is well-suited to measure accurately.

**Figure 5.**
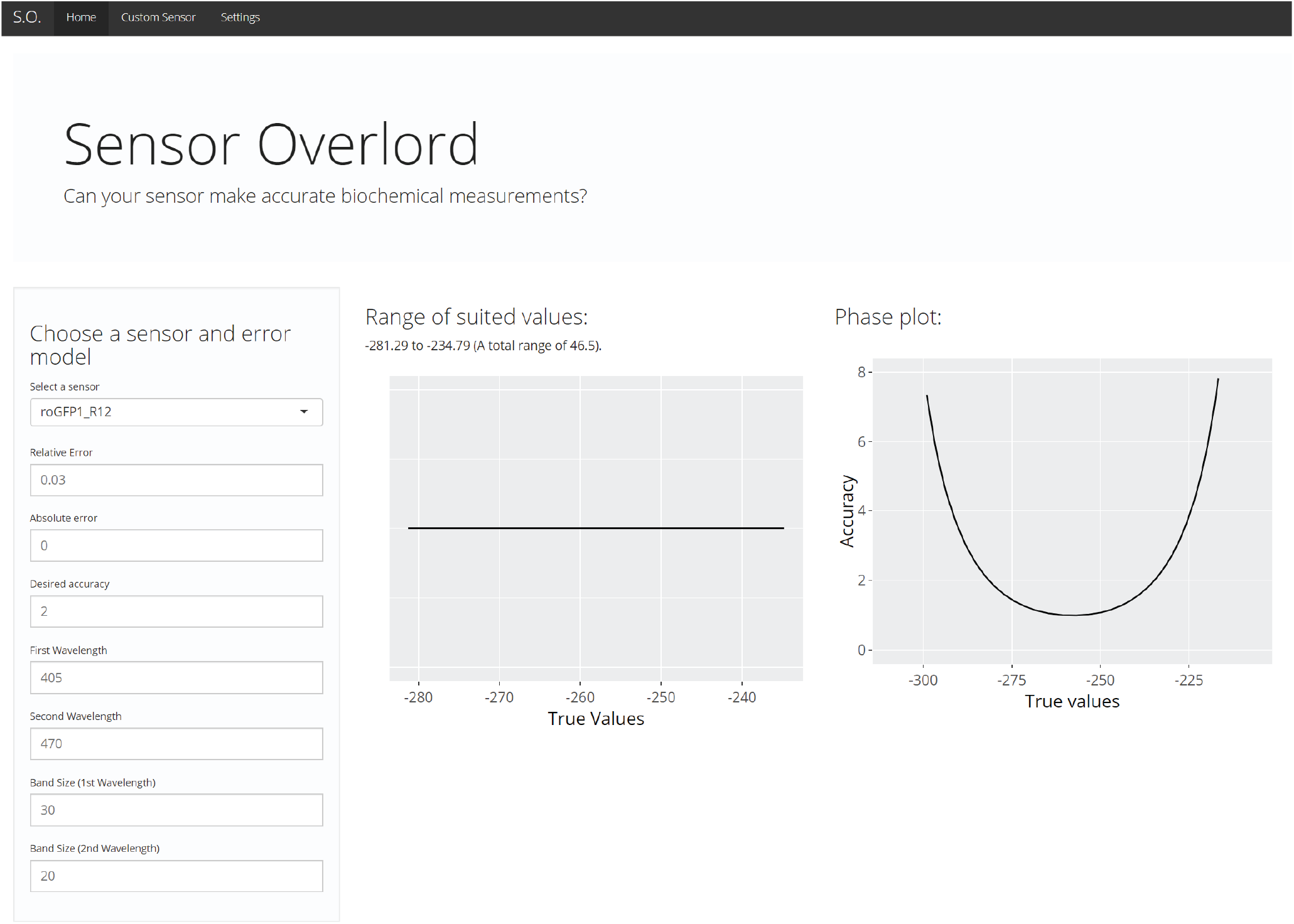
SensorOverlord web application. The SensorOverlord toolkit enables users to model how the range of input values that their biosensor is well-suited to measure accurately is constrained by the user’s fluorescence-ratio signal measurement precision and microscopy configuration.

## Discussion

The SensorOverlord toolkit enables users to predict the accuracy of concentrations and chemical potentials derived from fluorescence ratio measurements with two-state biosensors. This tool enables users to select biosensors predicted to be most accurate for measuring specific ranges of biochemical values. The SensorOverlord also enables users to quantify the extent to which increasing the precision of their fluorescence-ratio measurements would increase the predicted accuracy of their biochemical measurements with an individual biosensor. Therefore, this tool can be used to quantify the accuracy gains resulting from improving experimental practices, and from refining image acquisition, registration, and analysis methods.

A wide variety of factors can influence the precision of fluorescence-ratio measurement. In our experience, the degree of immobilization of live specimens during image acquisition can influence the precision of fluorescence-ratio measurements by a factor of three, leading to large differences in the predicted accuracy of biochemical measurements. The SensorOverlord enables researchers to disclose the predicted accuracy of the concentrations and chemical potentials that they measure, simply by reporting the precision of their fluorescence-ratio measurements—similar to how manufacturers use tolerance ratings to disclose how often the quality of their products is expected to deviate from a standard. The broader scientific community may, in turn, adopt appropriate maximum tolerable inaccuracy standards for specific biochemical measurements.

We hope that the SensorOverlord motivates the development of new biosensors, microscopy techniques, and image-analysis methods, by enabling biosensor developers and users to quantify the accuracy gains that would result from modifying the biochemical and spectral properties of their biosensors and from increasing the precision of their fluorescence-ratio measurements.

## Materials and Methods

### Code availability

Mathematical modeling was performed in the R language and environment for statistical computing (v3.6.0)^26^. The web application and associated visualizations were developed with the R packages ggplot2 (v3.1.1)^27^ and Shiny (v1.3.2)^28^, respectively. Source code for the SensorOverlord is available at https://github.com/julianstanley/SensorOverlord.

### Statistical analysis

All statistical analyses were performed in JMP (SAS). We tested for differences in the average *R* among groups using ANOVA. We used the Tukey HSD post-hoc test to determine which pairs of groups in the sample differ, in cases where more than two groups were compared. We used least-squares regression to quantify the dependency on *R* of the absolute error in *R* and the absolute relative error in *R*.

## Acknowledgements

We thank Jeff Bouffard, Jennifer Whangbo, Marianne Konikoff, Jodie Schiffer, Frank Servello, and Yuyan Xu for critical reading and detailed comments on our manuscript. We benefitted from discussions with members of Javier Apfeld’s and Erin Cram’s labs. Some strains were provided by the CGC, which is funded by NIH Office of Research Infrastructure Programs (P40 OD010440). The research was supported by a Northeastern TIER1 award and a National Science Foundation CAREER grant (1750065) to J.A.

## Author Contributions

J.A.S. and J.A. conceived and designed the analysis framework. J.A.S. designed and implemented the SensorOverlord software. S.B.J. analyzed fluorescence-ratio measurement precision. J.A.S. and J.A. analyzed data, interpreted results and wrote the manuscript.

