## Supplementary Information for "The SensorOverlord predicts the accuracy of measurements with ratiometric biosensors"

**Supplementary Note 1**

In this note we derive the map from roGFP-biosensor fluorescence-ratio ($R$) measurements into glutathione redox potential ($E_{GSH}$) values. We (1.1) review the general chemistry of roGFP biosensors; (1.2) derive the map between $R$ and $F_{ox}$; and (1.3) derive the map between $R$ and $E_{GSH}$.

**(1.1) General chemistry of roGFP biosensors**

roGFP biosensors include two cysteines whose thiol groups can form a reversible intramolecular disulfide bond [1]. Both reduced ($roGFP_{red}$) and oxidized ($roGFP_{ox}$) species exhibit green fluorescence but differ in the extent to which they fluoresce upon excitation at different wavelengths. Because these biosensors are derived from *Aequorea Victoria* green fluorescent protein (GFP), they exhibit two peaks in their excitation spectra, centered near 410 nm (A-band) and 470 nm (B-band), corresponding to the protonated and deprotonated forms of the biosensor's chromophore [2, 3]. The degree of chromophore protonation is influenced by the oxidation state of the two roGFP cysteines [4, 5]. As a result, the relative magnitude of these two excitation peaks differs between reduced and oxidized biosensor species. This means that when an ensemble of biosensors is excited at a given wavelength, the magnitude of their green-fluorescent emission depends not only on the number of biosensors but also on the fraction of biosensors in each oxidation state.

The fraction of oxidized biosensors ($F_{ox}$) provides a chemically interpretable description of the biosensor's oxidation. This fraction cannot be derived from their green-fluorescence at a single wavelength, when the biosensor concentration is not known. However, it is possible to calculate $F_{ox}$ if one records the ratio of the biosensor's green fluorescent emission at two different wavelengths ($R$) [1, 6], as we describe in section (1.2).

The tendency of $roGFP_{ox}$ to acquire electrons and thereby become reduced into $roGFP_{red}$ is quantified by the half-cell reduction potential of the $roGFP_{red}/roGFP_{ox}$ couple, $E_{roGFP}$. This redox potential is given by the Nernst equation:

$$E_{roGFP} = E^{\circ\prime}_{roGFP} - \frac{R_{gas}T}{2F}\ln\left(\frac{[roGFP_{red}]}{[roGFP_{ox}]}\right) \tag{1}$$

where $R_{gas}$ is the gas constant, $F$ is the Faraday constant, $T$ is the absolute temperature, and $E^{\circ\prime}_{roGFP}$ is the standard midpoint potential of the biosensor, which is -265 mV for roGFP1-R12 [7].

In section (1.3), we describe how one can calculate the biosensor's redox potential $E_{\text{roGFP}}$ given $F_{ox}$.

The oxidation and reduction of the roGFP-family of biosensors is controlled specifically by the glutathione/glutathione disulfide redox couple, in a manner catalyzed by the enzyme glutaredoxin [1, 8, 9]. The glutathione redox couple can be thought of as a broker that mediates the indirect effects of oxidants and reductants on the thiol-disulfide balance of many proteins, including roGFPs [6]. In *C. elegans* feeding muscles, roGFP1-R12 oxidation and reduction reactions in the cytosol exhibit fast kinetics *in vivo* [6]. As a result, the glutathione and roGFP redox couples are in equilibrium; that is, the redox potential of the biosensor's couple $E_{roGFP}$ is equal to the redox potential of the glutathione couple $E_{GSH}$ [6]. Measuring $E_{GSH}$ instead of $R$ enables us to make predictions about the oxidation state of the network of cysteines trading electrons with glutathione [6].

### (1.2) Derivation of the map between $R$ and $F_{ox}$

<u>(1.2.1) Dual-excitation single-emission biosensors</u>

Assume that a fluorescent biosensor has two states, $A$ and $B$, and that an individual biosensor's emission is state-dependent. Each biosensor in state $A$ emits an intensity $i_{\lambda,A}$ and each biosensor in state $B$ emits an intensity $i_{\lambda,B}$, where $\lambda$ represents that biosensor's <u>excitation wavelength</u>.

If an ensemble of $N_T$ biosensors were all in state A, the total observed intensity $I_{\lambda,A}$ would be:

$$I_{\lambda,A} = N_T\, i_{\lambda,A} \tag{2}$$

Similarly, if an ensemble of $N_T$ biosensors were all in state B, the total observed intensity $I_{\lambda,B}$ would be:

$$I_{\lambda,B} = N_T i_{\lambda,B} \tag{3}$$

In ensembles with $N_A$ biosensors in state $A$ and $N_B$ biosensors in state $B$, the total observed intensity $I_\lambda$ will be the sum of the intensities contributed by biosensors in each state:

$$I_\lambda = \frac{N_A}{N_T} I_{\lambda,A} + \frac{N_B}{N_T} I_{\lambda,B} \tag{4}$$

Since the biosensor can only be in one of two states, $N_A + N_B = N_T$, where $N_T$ is the total number of biosensors. Therefore:

$$I_\lambda = \frac{N_A}{N_T} I_{\lambda,A} + \left[1 - \frac{N_A}{N_T}\right] I_{\lambda,B} \tag{5}$$

We can express the probability of a biosensor being in state $A$ as $P(A) = N_A/N_T$ and the probability of a biosensor being in state $B$ as $P(B) = N_B/N_T = 1 - P(A)$. In a mixed population of biosensors, the total observed fluorescence intensity will be the weighted average of the intensities expected for populations of biosensors in each state:

$$I_\lambda = P(A)\, I_{\lambda,A} + [1 - P(A)]\, I_{\lambda,B} \tag{6}$$

By taking a ratiometric measurement $R$:

$$R = \frac{I_{\lambda_1}}{I_{\lambda_2}} = \frac{\frac{N_A}{N_T}\, I_{\lambda_1,A} + \left[1 - \frac{N_A}{N_T}\right] I_{\lambda_1,B}}{\frac{N_A}{N_T}\, I_{\lambda_2,A} + \left[1 - \frac{N_A}{N_T}\right] I_{\lambda_2,B}} \tag{7}$$

which simplifies to, in terms of $P(A)$:

$$P(A) = \frac{I_{\lambda_2} I_{\lambda_1,B} - I_{\lambda_1} I_{\lambda_{2,B}}}{I_{\lambda_1} I_{\lambda_{2,A}} - I_{\lambda_1} I_{\lambda_2,B} - I_{\lambda_2} I_{\lambda_1,A} + I_{\lambda_2} I_{\lambda_1,B}} \tag{8}$$

Now, assume that the maximum possible $R$ measurement occurs when all biosensors are in state $A$ (that is, when $P(A) = 1$) and that the minimum possible $R$ occurs when all biosensors are in state $B$ (that is, when $P(A) = 0$). We can thereby define three constants:

$$R_A = \frac{I_{\lambda_1,A}}{I_{\lambda_2,A}} \tag{9}$$

$$R_B = \frac{I_{\lambda_1,B}}{I_{\lambda_2,B}} \tag{10}$$

$$\delta_{\lambda_2} = \frac{I_{\lambda_2,A}}{I_{\lambda_2,B}} \tag{11}$$

And express $P(A)$ in terms of $R$ and those three constants:

$$P(A) = \frac{R - R_B}{R - R_B + \delta_{\lambda_2}(R_A - R)} \tag{12}$$

Then, if biosensors in state $A$ are oxidized, biosensors in state *B* are reduced, $\lambda_1 = 410\ nm$, and $\lambda_2 = 470\ nm$, Equation (12) becomes the map between $R$ and $F_{ox}$ (Equation 16):

$$R_A = R_{ox} = \frac{I_{410,ox}}{I_{470,ox}} \tag{13}$$

$$R_B = R_{red} = \frac{I_{410,red}}{I_{470,red}} \tag{14}$$

$$\delta_{\lambda_2} = \delta_{470} = \frac{I_{470,ox}}{I_{470,red}} \tag{15}$$

$$P(A) = F_{ox} = \frac{R - R_{red}}{R - R_{red} + \delta_{470}(R_{ox} - R)} \tag{16}$$

where $R_{red}$ is the ratiometric emission of an ensemble of reduced biosensors, $R_{ox}$ is the ratiometric emission of an ensemble of oxidized biosensors, and $\delta_{470}$ is the biosensor's dynamic range in the second wavelength, where the dynamic range $DR = \delta_{410}/\delta_{470} = R_{ox}/R_{red}$.

(1.2.2) Dual-emission single-excitation biosensors

Here, as with dual-excitation single-emission biosensors (see section 1.2.1) we assume that the fluorescent biosensor has two states, $A$ and $B$, and that an individual biosensor's emission is state-dependent. In this case, however, each biosensor in state $A$ emits an intensity $i_{\lambda,A}$ and each biosensor in state $B$ emits an intensity $i_{\lambda,B}$, where $\lambda$ represents that biosensor's emission wave-length.

(1.2.3) Dual-excitation and dual-emission biosensors

Here, as before (see sections 1.2.1 and 1.2.2) we assume that the fluorescent biosensor has two states, $A$ and $B$, and that an individual biosensor's emission is state-dependent. In this case, however, each biosensor in state $A$ emits an intensity $i_{\lambda,A}$ and each biosensor in state $B$ emits an intensity $i_{\lambda,B}$, where $\lambda$ represents that biosensor's excitation- and emission-wavelength pair. This class of biosensors could, in principle, be built by linking two fluorescent proteins, each with well-separated maxima of fluorescence excitation and emission, with the fluorescence of the former being input-sensitive and the fluorescence of the latter being input-insensitive.

#### (1.3) Derivation of the map between $R$ and $E_{GSH}$

The reduction potential of a two-state redox biosensor is given by the Nernst equation:

$$E_{redox\ biosensor} = E^{\circ\prime}_{redox\ biosensor} - \frac{R_{gas}T}{2F}\ln\left(\frac{[roGFP_{red}]}{[roGFP_{ox}]}\right) \tag{17}$$

In *in vivo* experiments, the concentration of the biosensor is generally not known *a priori*. The concentrations of reduced and oxidized biosensor species, however, are constrained by mass balance:

$$[roGFP] = [roGFP_{red}] + [roGFP_{ox}] \tag{18}$$

Given this constraint, we can express $E_{biosensor}$ in terms of $F_{ox}$:

$$[roGFP] = \frac{N_T}{Volume} = \frac{N_{red}}{Volume} + \frac{N_{ox}}{Volume} \tag{19}$$

so

$$\frac{[roGFP_{red}]}{[roGFP_{ox}]} = \frac{N_T - N_{ox}}{N_{ox}} = \frac{1 - N_{ox}/N_T}{N_{ox}/N_T} = \frac{1 - F_{ox}}{F_{ox}} \tag{20}$$

and

$$E_{redox\ biosensor} = E^{\circ\prime}_{redox\ biosensor} - \frac{R_{gas}T}{2F}\ln\left(\frac{1 - F_{ox}}{F_{ox}}\right) \tag{21}$$

For roGFP biosensors, then:

$$E_{roGFP} = E^{\circ\prime}_{roGFP} - \frac{R_{gas}T}{2F}\ln\left(\frac{1 - F_{ox}}{F_{ox}}\right) \tag{22}$$

For two-state ratiometric biosensors, we can use Equation (16) to write $F_{ox}$ in terms of the fluorescence ratio $R$, and this expression simplifies to Equation (23):

$$E_{roGFP} = E^{\circ\prime}_{roGFP} - \frac{R_{gas}T}{2F}\ln\left(\delta_{\lambda_2}\frac{R_{ox} - R}{R - R_{red}}\right) \tag{23}$$

For roGFP biosensors in equilibrium with the glutathione redox couple, then:

$$E_{GSH} = E_{roGFP} = E^{\circ\prime}_{roGFP} - \frac{R_{gas}T}{2F}\ln\left(\delta_{\lambda_2}\frac{R_{ox} - R}{R - R_{red}}\right) \tag{24}$$

**Supplementary Note 2**

In Supplementary Note 1 we derived the map from roGFP-biosensor fluorescence-ratio ($R$) measurements into the fraction of oxidized biosensors ($F_{ox}$). Knowing $F_{ox}$ enables calculation of the biosensor's redox potential ($E_{roGFP}$), which equals the glutathione redox potential ($E_{GSH}$) under equilibrium conditions. In this note we determine (2.1) the sensitivity of the fraction of oxidized biosensors to errors in $R$; and (2.2) the sensitivity of $E_{GSH}$ values to errors in $R$.

**(2.1) Sensitivity of the fraction of oxidized biosensors to biosensor fluorescence-ratio**

For small errors in $R$, the sensitivity of $F_{ox}$ to changes in $R$ is quantified by its partial derivative with respect to $R$:

$$F_{ox} = \frac{R - R_{red}}{R - R_{red} + \delta_{\lambda_2}(R_{ox} - R)} \tag{1}$$

$$\frac{\partial F_{ox}}{\partial R} = \frac{\delta_{\lambda_2}(R_{ox} - R_{red})}{\left(R(\delta_{\lambda_2} - 1) - R_{ox}\delta_{\lambda_2} + R_{red}\right)^2} \tag{2}$$

where $R_{red}$ is the ratiometric emission of an ensemble of reduced biosensors, $R_{ox}$ is the ratiometric emission of an ensemble of oxidized biosensors, and $\delta_{\lambda_2}$ is the biosensor dynamic range in the second excitation band (*e.g.* in our experiments in *C. elegans*, where we excite roGFP1-R12 with bands of light $\lambda_1$and $\lambda_2$ centered at 410 nm and 470 nm, respectively, $\delta_{\lambda_2}$ corresponds to the dynamic range of the 470 nm excitation band [6]).

The sensitivity of $F_{ox}$ with respect to $R$ depends on $\delta_{\lambda_2}$:

- When $\delta_{\lambda_2}$ is 1, the map from $R$ to $F_{ox}$ is linear (Figure S2-1a, black line) and, therefore, $\partial F_{ox}/\partial R$ is equal to $1/(R_{ox} - R_{red})$ throughout the range of ratio values (Figure S2-1b, black line). We note this corresponds to the special case where reduced and oxidized biosensor forms have the same fluorescence upon excitation in the second wavelength (*i.e.* $\lambda_2$ is an isosbestic point); as a result, the biosensor's fluorescence upon excitation with this wavelength does not contribute to the overall dynamic range of the biosensor fluorescence ratio (*DR*), equal to $\delta_{\lambda_1}/\delta_{\lambda_2}$.
- When $\delta_{\lambda_2}$ is less than 1, the map from $R$ to $F_{ox}$ is non-linear (Figure S2-1a, red curves). Here, the sensitivity of $F_{ox}$ with respect to $R$ is highest when $R$ equals its lowest value, $R_{red}$, and

decreases monotonically as $R$ approaches its maximum value, $R_{ox}$ (Figure S2-1b, red curves). We note that in this case the biosensor's fluorescence upon excitation with each of $\lambda_1$and $\lambda_2$ contributes to the overall dynamic range of the biosensor's fluorescence ratio.

- When $\delta_{\lambda_2}$ is greater than 1, the map from $R$ to $F_{ox}$ is also non-linear (Figure S2-1a, blue curves). Here, the sensitivity of $F_{ox}$ with respect to $R$ is lowest when $R$ equals its lowest value, $R_{red}$, and increases monotonically as $R$ approaches its maximum value, $R_{ox}$ (Figure S2-1a, blue curves). We note that in this case $\lambda_1$ increases the overall dynamic range of the biosensor's fluorescence ratio, but $\lambda_2$ decreases it.

**(2.2) Sensitivity of the biosensor's redox potential to biosensor fluorescence-ratio**

For small errors in $R$, the sensitivity of $E_{GSH}$ to changes in $R$ is quantified by its partial derivative with respect to $R$:

$$E_{roGFP} = E^{\circ\prime}_{roGFP} - \frac{R_{gas}T}{2F} \ln\,(\delta_{\lambda_2} \frac{R_{ox} - R}{R - R_{red}}) \qquad (3)$$

$$\frac{\partial E_{roGFP}}{\partial R} = \frac{R_{gas}T}{2F} * \frac{R_{ox} - R_{red}}{(R_{ox} \; - R)(R - R_{red})} \qquad (4)$$

The map from $R$ to $E_{GSH}$ depends on both $R$ and $\delta_{\lambda_2}$ (Figure S2-1c). The sensitivity of $E_{GSH}$ to $R$ varies strongly with $R$ (Figure S2-1d). We note that $\partial E_{roGFP}/\partial R$ is at its lowest value when $R$ is halfway between its upper and lower bounds, $R_{ox}$ and $R_{red}$, but increases rapidly as $R$ approaches either one of those bounds (Figure S2-2b).

The sensitivity of $E_{roGFP}$ to $R$ is independent of $\delta_{\lambda_2}$ (Figure S2-1d). This is in contrast with the sensitivity of $F_{ox}$ to $R$, which has a complex dependence on $\delta_{\lambda_2}$ (Figure S2-1b).

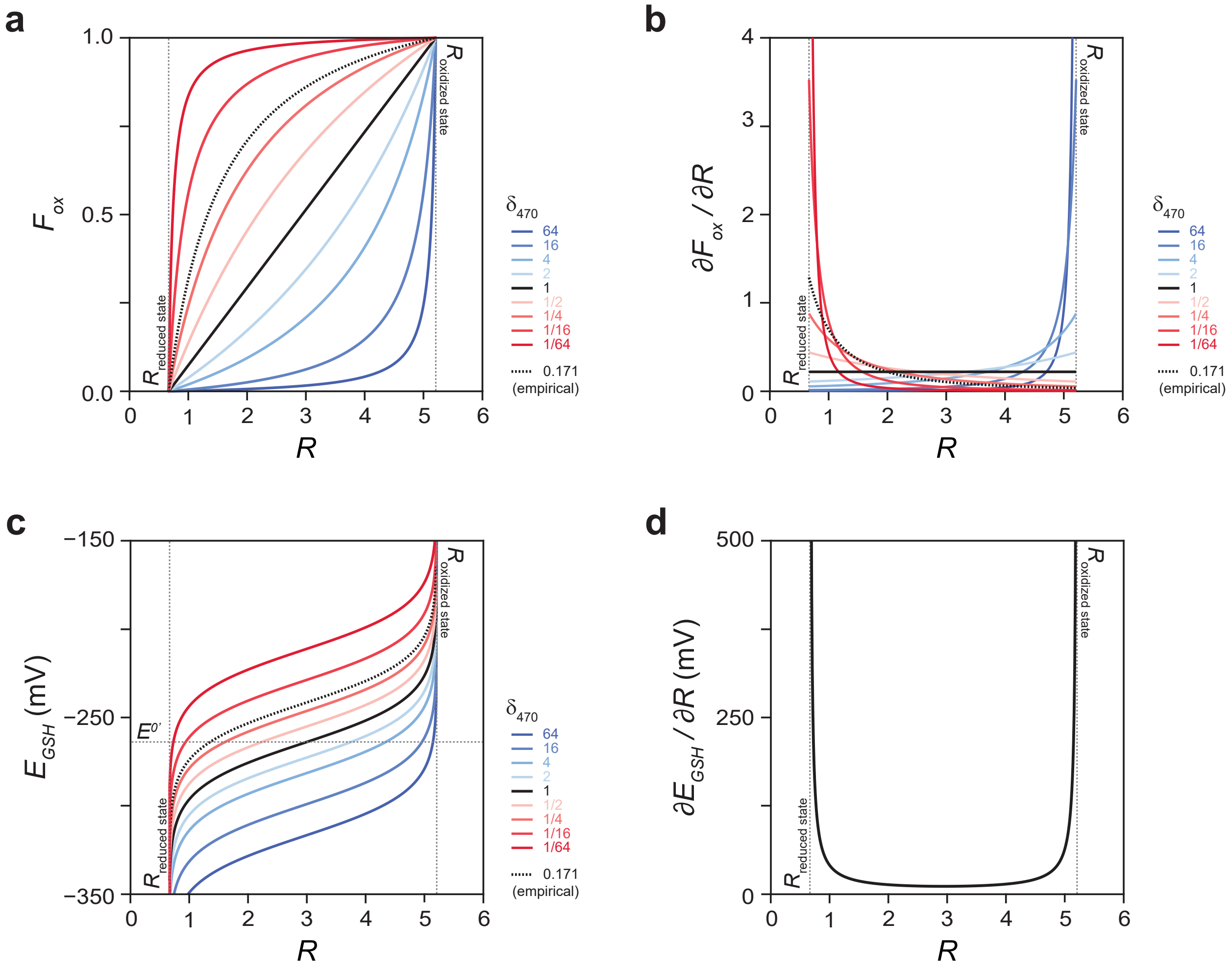


**Figure S2-1: Sensitivity of the fraction of oxidized biosensors and glutathione redox potential $E_{GSH}$ values to errors in fluorescence-ratio $R$**

**a.** The conversion map from the biosensor fluorescence-ratio $R$ to the fraction of oxidized biosensors $F_{ox}$ depends on the ratiometric emission of the ensemble of reduced biosensors $R_{red}$, the ratiometric emission of the ensemble of oxidized biosensors $R_{ox}$, and the biosensor dynamic range in the second excitation band $\delta_{\lambda_2}$.

**b.** The sensitivity of $F_{ox}$ to changes in $R$ has a complex dependence on both $R$ and $\delta_{\lambda_2}$.

**c.** The conversion map from $R$ to $E_{GSH}$ depends on both $R$ and $\delta_{\lambda_2}$.

**d.** The sensitivity of $E_{GSH}$ to changes in $R$ depends on $R$ but is independent of $\delta_{\lambda_2}$.

In panels **a**, **b***,* and **c**, the dotted curve was calculated using $R_{red}$ $R_{ox}$, and $\delta_{\lambda_2}$values determined empirically for our experimental setup using live *C. elegans* expressing roGFP1-R12 [6].

**Supplementary Note 3**

In this note, we quantify empirical errors in $R$ in live *C. elegans* expressing the roGFP1-R12 biosensor in the cytosol of the muscles of the pharynx, the feeding organ. This retrospective analysis of thousands of fluorescence-ratio images showed that (3.1) our errors in $R$ were relative—that is, $R_{obs} = R_{true} \times (1 + error)$—and were invariant over the range of all possible $R$ values; and that (3.2) these relative errors vary between experiments.

**(3.1) The relative error in *R* is independent of *R***

To determine whether the error in our $R$ measurements varies with $R$, we quantified $R$ errors in experiments in which the fraction of oxidized biosensors was manipulated by exposing *C. elegans* worms to different chemicals [6]. Each experiment consisted of a collection of time series where $R_{410/470}$ was measured longitudinally in individual worms first under normal conditions ("baseline") and then in response to treatment with 50 mM diamide (a thiol-specific oxidant). These animals where then either treated with 100 mM dithiothreitol (DTT, a reducing agent) or returned to normal conditions ("recovery").

The biosensor's green fluorescence ratio ($R_{obs}$) is the ratio ($R_{410/470}$) of the observed fluorescence intensities on excitation with 410 nm and 470 nm light ($I_{410}$ and $I_{470}$, respectively) captured via sequential imaging. Under baseline conditions $R_{obs}$ was in a steady state or changed slowly over the measurement interval of ten minutes [6] (Figure S3-1a). Treatment with 50 mM diamide caused a new steady state in $R$ consistent with maximal oxidation of the biosensor [6] (Figure S3-1a). Transferring animals from 50 mM diamide back to normal conditions resulted in spontaneous recovery from maximal oxidation and a new steady state in $R$ near baseline values [6] (Figure S3-1a). To estimate $R_{true}$, we fitted linearly each $R_{obs}$ time series within a 10-minute interval corresponding to the baseline or the end of each treatment. We defined the error in $R$ as the difference between $R_{obs}$ and $R_{true}$ and defined the relative error in $R$ as the error in $R$ divided by $R_{true}$. The error in *R* increased in response to diamide treatment and decreased upon recovery from diamide (Figure S3-1b). In contrast, the relative error in $R$ was unaffected by diamide treatment or recovery (Figure S3-1c). These findings suggested the relative error in $R$ is independent of $R$.

To estimate the error in $R$ over a broader range of values, we performed the same analysis for the collection of time series in which animals were transferred first to diamide and then to DTT. We found the same pattern: diamide treatment increased the error in $R$ and subsequent DTT treatment reduced it (Figure S3-2a); however, these treatments had only very small effects on the relative error in $R$ (Figure S3-2b). We then re-analyzed both collections of $R$ time series, but this time computed the errors in the observed $R_{470/410}$ values using the same approach as for the errors in $R_{410/470}$. Because $R_{470/410}$ is the inverse of $R_{410/470}$, this approach enabled us to eliminate many gaps in $R$ coverage. The observed fluorescence ratios and their inverses varied over a wide range of values and exhibited large difference in their absolute errors (Figure S3-2a,c); however, their relative errors exhibited only very small differences (Figure S3-2b,d). These results confirm that in our experiments, the relative error in $R$ was independent of $R$.

#### (3.2) The relative error in *R* can vary between experiments

To determine whether the relative error in $R$ varies between experiments, we followed the same approach as in (3.1) and analyzed three additional collections of time series of animals under baseline conditions. We found that the relative error in $R$ varied between the baselines of the five experiments by up to three-fold (Figure S3-3). Differences in the proportion of animals moving during imaging account for most of the variation in the relative error in *R* across experiments (S.B.J, J.A.S, and J.A., manuscript in preparation).

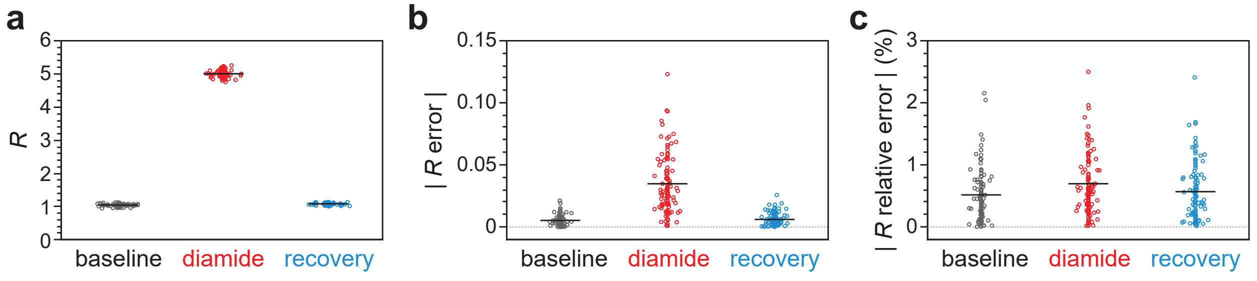


**Figure S3-1: The error in $R$ is relative**

**a.** $R_{410/470}$ measurements in *C. elegans* worms expressing roGFP1-R12 in feeding muscles under normal conditions ("baseline"), in response to treatment with 50 mM diamide, and upon return to normal conditions ("recovery").

**b.** The absolute value of the error in $R$ varies between baseline, diamide treatment, and recovery ($P < 0.0001$, ANOVA).

**c.** The absolute value of the relative error in $R$ does not vary between baseline, diamide treatment, and recovery conditions ($P > 0.05$, ANOVA).

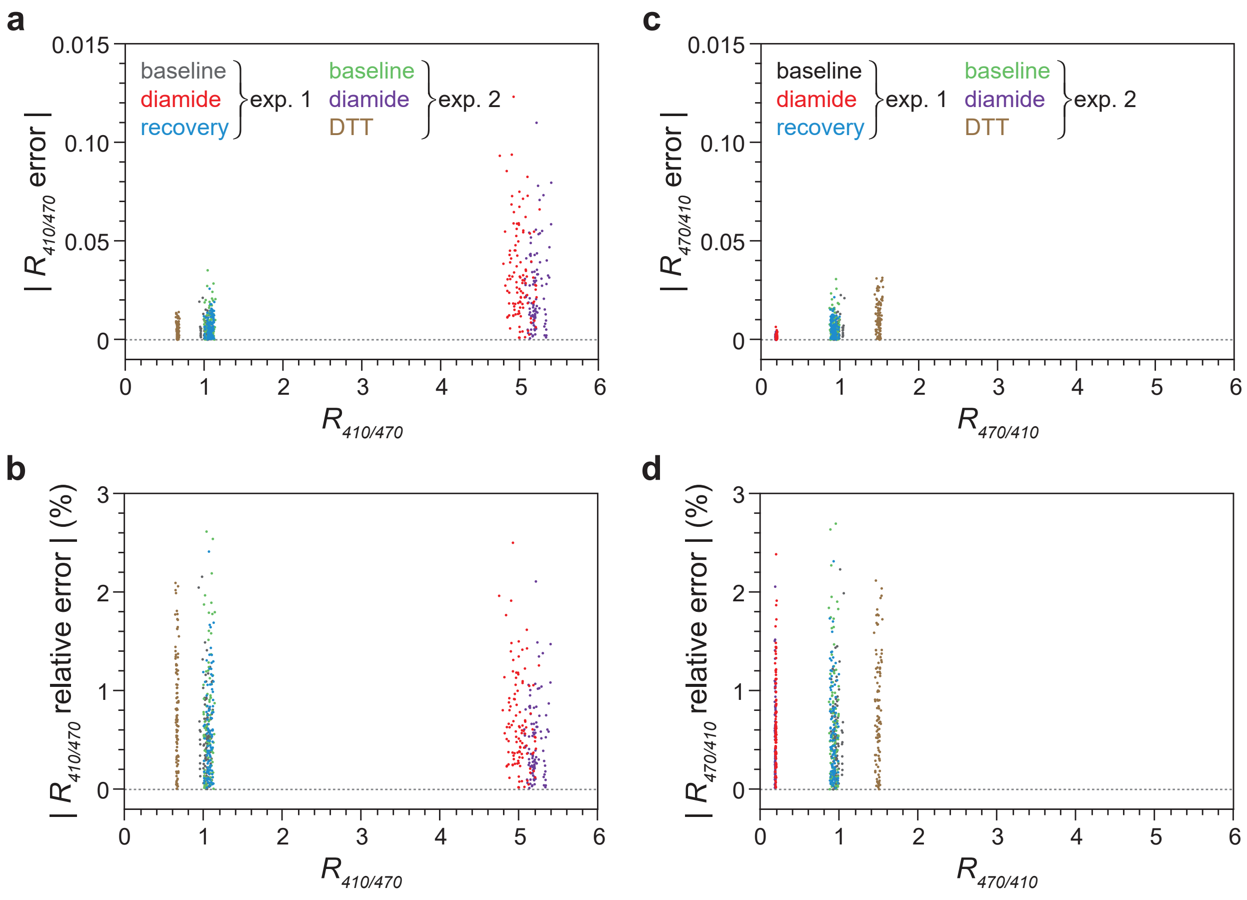


**Figure S3-2: The relative error in $R$ is independent of $R$ over a broad range of $R$ values**

**a.** The absolute value of the error in $R_{410/470}$ varies with $R_{410/470}$ ($P$ < 0.0001, linear regression).

**b.** The absolute value of the relative error in $R_{410/470}$ does not vary with $R_{410/470}$ ($P$ > 0.05, linear regression).

**c.** The absolute value of the error in $R_{470/410}$ varies with $R_{470/410}$ ($P$ < 0.0001, linear regression).

**d.** The absolute value of the relative error in $R_{470/410}$ does not vary with $R_{470/410}$ ($P$ > 0.05, linear regression).

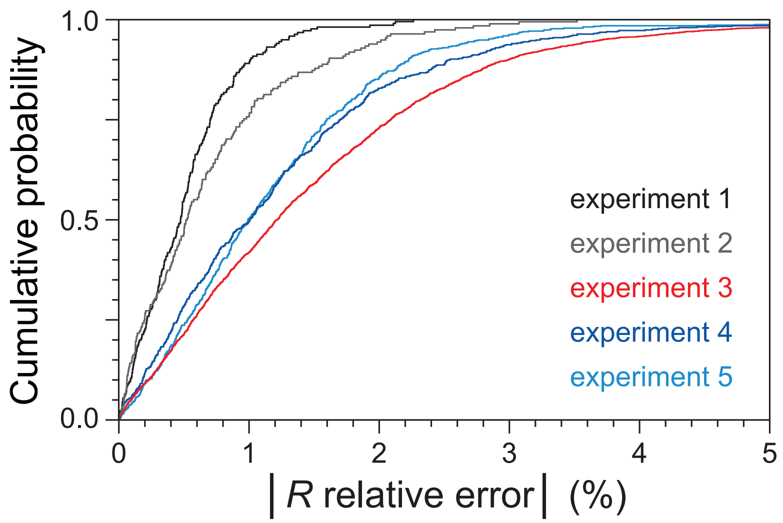


**Figure S3-3: The relative error in $R$ can vary between experiments**

Cumulative distributions of the absolute value of the relative error in $R_{410/470}$ in five experiments consisting of baseline measurements of $R_{410/470}$. The absolute value of the relative error in $R_{410/470}$ varied between pairs of experiments ($P < 0.0001$, Turkey HSD test), except for experiments 1 and 2 and experiments 4 and 5, which did not exhibit differences ($P > 0.05$, Turkey HSD test).

**Supplementary Note 4**

In this note, we build on the framework described in Supplementary Note 1 to model how the biochemical and biophysical properties of roGFP biosensors affect the relationship between the empirical precision of our fluorescence-ratio measurements and the accuracy of individual $E_{GSH}$ observations: we (4.1) present the overall approach; (4.2) show how $E_{GSH}$ measurement inaccuracy is sensitive to the biochemical and biophysical variables that map $R$ to $E_{GSH}$; and (4.3) extend this analysis to other types of ratiometric two-state biosensors.

**(4.1) Overall approach**

In Supplementary Note 1.3 we derived the function mapping the ratiometric emission ($R$) of dual-excitation single emission roGFP biosensors and the reduction potential of the glutathione-glutathione disulfide couple $E_{GSH}$:

$$E_{GSH} = E_{roGFP} = E^{\circ\prime}_{roGFP} - \frac{R_{gas}T}{2F}\ln\left(\delta_{\lambda_2}\frac{R_{ox} - R}{R - R_{red}}\right) \quad (1)$$

The function mapping $R$ to $E_{GSH}$ depends on physical, biochemical, and biophysical parameters. These include (a) physical constants and variables: the gas constant $R_{gas}$, the Faraday constant $F$, and the absolute temperature $T$; (b) biochemical variables: the standard midpoint potential of the biosensor $E^{\circ\prime}_{roGFP}$; and (c) biophysical variables: the ratiometric emission of an ensemble of reduced biosensors $R_{red}$, the ratiometric emission of an ensemble of oxidized biosensors $R_{ox}$, and the biosensor's dynamic range in the second excitation wavelength $\delta_{\lambda_2}$.

In the main text, we modeled how the empirical error in $R$ measurement influences the accuracy of individual $E_{GSH}$ measurements (Figure 1g). In this note, we model how that relationship is influenced by each of the biochemical and biophysical variables mapping $R$ to $E_{GSH}$.

**(4.2) Sensitivity of $E_{GSH}$ measurement inaccuracy to the biochemical and biophysical variables that map $R$ to $E_{GSH}$**

In this analysis, we changed the values of individual parameters of the function mapping $R$ to $E_{GSH}$ while holding other parameters constant. We then determined how the inaccuracy of our $E_{GSH}$ measurements would be affected by a 2.8%-fold error in $R$. This relative error corresponds to the 0.025 and 0.975 quartiles of our empirical distribution of relative errors in $R$ (the median relative error in $R$ was zero) (Figure 1f). Thus, the $E_{GSH}$ inaccuracy we calculated is the maximum absolute difference between $E_{True}$ and either the upper or lower bounds of the interval encompassing 95% of the $E_{Observed}$ values.

As a starting point we considered a hypothetical roGFP biosensor with $E°'$ = -250 mV. Other parameters were set as follows: $R_{red}$ = 1, $R_{ox}$ = 5, and $\delta_{\lambda_2}$ = 1. We note the constraints between certain parameters given by the biosensor's dynamic range $DR$:

$$DR = \frac{R_{ox}}{R_{red}} = \frac{\delta_{\lambda_1}}{\delta_{\lambda_2}} \tag{2}$$

Changing the biosensor's midpoint potential shifts the x-axis of the map between the true value of $E_{GSH}$ and $E_{GSH}$ inaccuracy (Figure S4-1a). The map between $E_{GSH} - E°'$ and $E_{GSH}$ inaccuracy is unaffected by the value of $E°'$.

Changing the biosensor's $\delta_{\lambda_2}$ while holding $DR$ constant also shifted the x-axis of the map between the true value of $E_{GSH}$ and $E_{GSH}$ inaccuracy (Figure S4-1b). An $x$-fold change in $\delta_{\lambda_2}$ while holding $DR$ constant shifts the x-axis of that map by $-\frac{R_{gas}T}{2F}\ln x$.

Changing the biosensor's $\delta_{\lambda_1}$ without the constraint of holding $DR$ constant had a more complex effect (Figure S4-1c). Increasing $\delta_{\lambda_1}$ (and thereby $DR$) always lowered $E_{GSH}$ inaccuracy. In addition, it decreased the $E_{GSH}$ value that maps to the lowest $E_{GSH}$ inaccuracy. Changing $\delta_{\lambda_2}$ without the constraint of holding $DR$ constant had a similar effect to changing $\delta_{\lambda_1}$ (Figure S4-1d). At a given $DR$ value, the curves mapping $E_{GSH}$ to $E_{GSH}$ inaccuracy are identical if one changed either $\delta_{\lambda_1}$ or $\delta_{\lambda_2}$ except for a shift in the x-axis (due to changing $\delta_{\lambda_2}$at constant $DR$, described above).

In summary, our sensitivity analysis indicates that the optimal strategy for minimizing $E_{GSH}$ inaccuracy with a given biosensor and error level in $R$ is to select excitation wavelengths that maximize the biosensor's dynamic range $DR$. Choosing a combination of wavelengths that includes a wavelength at the isosbestic point of a biosensor (*i.e.* where $\delta$ = 1) provides a linear map between $R$ and the biosensor's fraction oxidized $F_{ox}$ (Figure S2-1a); however, such wavelength combinations would always result in higher $E_{GSH}$ inaccuracy than combinations with a higher $DR$.

**(4.3) Generalization to other types of ratiometric biosensors**

In Supplementary Note 1.2, we showed that the function mapping $R$ to $E_{GSH}$ (equation 1 in this note) generalizes to dual-emission single-excitation biosensors, and to dual-excitation and dual-emission biosensors; and in Supplementary Note 7 we show that a function of the same general form applies to pH and ligand-binding biosensors. Therefore, the conclusions of the sensitivity analysis presented in this Supplementary Note apply to all these types of biosensors.

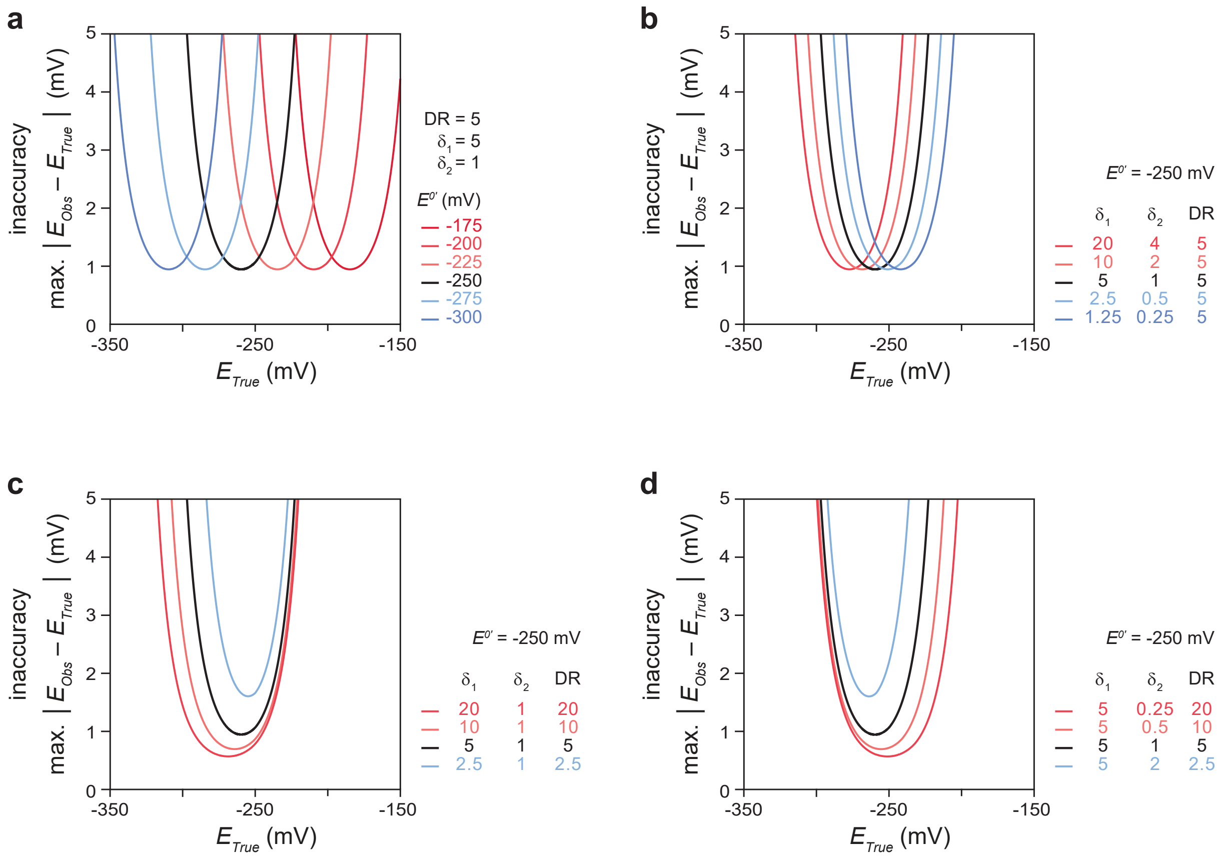


**Figure S4-1: Sensitivity of $E_{GSH}$ measurement inaccuracy to the biochemical and biophysical variables that map $R$ to $E_{GSH}$**

**a.** Sensitivity of $E_{GSH}$ measurement inaccuracy (the maximum absolute difference between $E_{True}$ and $E_{Observed}$) to the standard midpoint potential of the biosensor $E°'$, when other parameters mapping $R$ to $E_{GSH}$ are held constant.

**b.** Sensitivity of $E_{GSH}$ measurement inaccuracy to the biosensor's dynamic range in the first and second excitation wavelengths, $\delta_1$ and $\delta_2$, respectively, when holding constant the biosensor's dynamic range $DR$ and the biosensor's $E°'$.

**c.** Sensitivity of $E_{GSH}$ measurement inaccuracy to the biosensor's dynamic range in the first excitation wavelength $\delta_1$, when holding constant the biosensor's dynamic range in the second excitation wavelength $\delta_2$ and the biosensor's $E°'$.

**d.** Sensitivity of $E_{GSH}$ measurement inaccuracy to the biosensor's dynamic range in the second excitation wavelength $\delta_2$, when holding constant the biosensor's dynamic range in the first excitation wavelength $\delta_1$ and the biosensor's $E°'$.

In all panels, the relative error in $R$ was set to 2.8%-fold, which corresponds to the 0.025 and 0.975 quartiles of the empirical distribution of relative errors in $R$ in our experiments with roGFP1-R12 in the feeding muscles of *C. elegans* (Figure 1f).

**Supplementary Note 5**

In Supplementary Note 1 we derived the function mapping the ratiometric emission ($R$) of dual-excitation single emission roGFP biosensors and the reduction potential of the glutathione-glutathione disulfide couple $E_{GSH}$. That map depends on physical, biochemical, and biophysical parameters, which either quantify intrinsic properties of the biosensor or vary between microscope setups. Knowledge of these parameters enables modeling how the precision of $R$ measurements with a biosensor determines the range of $E_{GSH}$ values that is possible to measure with that biosensor at a given inaccuracy level. In this note, we (5.1) determine the minimal set of parameters and constraints between parameters needed to predict the influence of relative errors in $R$ on $E_{GSH}$ measurement error; (5.2) estimate the values of those parameters for eleven roGFP-based biosensors with known midpoint potentials and fluorescence spectra; and (5.3) determine the $E_{GSH}$ inaccuracy we would expect to observe had we measured $E_{GSH}$ in the feeding muscles of live *C. elegans* with each of those biosensors instead of roGFP-R12.

**(5.1) Which of the parameters mapping a biosensor's fluorescence ratio $R$ to $E_{GSH}$ are needed to model how relative errors in $R$ influence $E_{GSH}$ measurement error?**

<u>(5.1.1) What are the values of the parameters mapping a biosensor's fluorescence ratio $R$ to $E_{GSH}$?</u>

The function mapping the ratiometric emission ($R$) of dual-excitation single emission roGFP biosensors and the reduction potential of the glutathione-glutathione disulfide couple $E_{GSH}$, was derived in Supplementary Note 1.3:

$$E_{GSH} = E_{roGFP} = E^{\circ\prime}_{roGFP} - \frac{R_{gas}T}{2F}\ln\left(\delta_{\lambda_2}\frac{R_{ox}-R}{R-R_{red}}\right) \quad (1)$$

That map depends on physical, biochemical, and biophysical parameters. The physical parameters are generally known; they include the gas constant $R_{gas}$, the Faraday constant $F$, and the absolute temperature $T$. The biochemical parameter, the standard midpoint potential of the biosensor $E^{\circ\prime}_{roGFP}$, is often known from published reports. The biophysical parameters include the biosensor's dynamic range in the second excitation wavelength $\delta_{\lambda_2}$, the ratiometric emission of an ensemble of reduced biosensors $R_{red}$, and the ratiometric emission of an ensemble of oxidized biosensors $R_{ox}$. These three parameters can be determined empirically, as we have done previously in live *C. elegans* expressing roGFP1-R12 [6], but in most cases they are not known. The value of $\delta_{\lambda_2}$ can be determined from the fluorescence spectra of reduced and oxidized biosensors (section 5.1.2). These spectra also produce $R_{ox}$ and $R_{red}$ values (section 5.1.2). However, these values vary from one imaging set up to another. Because $R_{ox}$ and $R_{red}$ values are instrument specific, the biosensor's ratiometric emission $R$ collected in a microscope cannot be mapped to $E_{GSH}$ using the $R_{ox}$ and $R_{red}$ values obtained from a different instrument, such as spectrofluorometer used to collect the excitation spectra of reduced and oxidized biosensor species. The variation of $R_{ox}$ and $R_{red}$ across instruments, however, is constrained by the biophysical properties of the biosensor at the excitation wavelengths $\lambda_1$ and $\lambda_2$, such that the ratio of $R_{ox}$ to $R_{red}$ (the biosensor's overall dynamic range, $DR$) is constant. $DR$ can be determined from the fluorescence spectra of reduced and oxidized biosensors (section 5.1.2). Therefore, in order to map $R$ to $E_{GSH}$ for a given biosensor, either $R_{ox}$ or $R_{red}$ must be determined empirically in the same instrument where that biosensor's $R$ is measured.

Sometimes, all parameters mapping a biosensor's ratiometric emission in a given instrument to $E_{GSH}$ values are known, except for $R_{ox}$ and $R_{red}$. Even though in those cases it is not possible to map a biosensor's $R$ to $E_{GSH}$, it is still desirable to determine how relative errors in that biosensor's $R$ would influence $E_{GSH}$ measurement error. For example, such analysis may enable comparison of the expected $E_{GSH}$ inaccuracy of different biosensors under otherwise identical microscopy conditions. In section (5.1.3), we demonstrate that knowledge of the biosensor's overall dynamic range $DR$ is sufficient to model how relative errors in a biosensor's $R$ influence $E_{GSH}$ measurement error with that biosensor when the values of $R_{red}$ and $R_{ox}$ are not known.

<u>(5.1.2) Obtaining $\delta_{\lambda_2}$ and $R_{ox}/R_{red}$ from the excitation spectra of reduced and oxidized biosensor species</u>

Equation (1) can be reparametrized in terms of the biosensor's overall dynamic range $DR$:

$$DR = \frac{R_{ox}}{R_{red}} = \frac{\delta_{\lambda_1}}{\delta_{\lambda_2}} \tag{2}$$

$$E_{GSH} = E_{roGFP} = E^{\circ\prime}_{roGFP} - \frac{R_{gas}T}{2F} \ln\left(\delta_{\lambda_2} \frac{DR - R/R_{red}}{R/R_{red} - 1}\right) \tag{3}$$

where $\delta_\lambda$ is the ratio of the biosensor's emission when excited at $\lambda$ in the oxidized and reduced states:

$$\delta_\lambda = \frac{i_{\lambda,ox}}{i_{\lambda,red}} \tag{4}$$

$\delta_\lambda$ is, therefore, independent of the illumination power at $\lambda$ and should be invariant across instruments.

The overall dynamic range $DR$ of a dual-excitation biosensor is the ratio of the $\delta_\lambda$values at each excitation wavelength. $DR$ is also independent of the illumination power at $\lambda_1$ and $\lambda_2$, and should be invariant across instruments.

(5.1.3) Modeling how relative errors in $R$ influence $E_{GSH}$ measurement error when the values of $R_{red}$ and $R_{ox}$ are not known but their ratio is known

Unlike $DR$ and $\delta_\lambda$, $R_{ox}$ and $R_{red}$ are instrument-dependent and must be determined empirically for each imaging instrument. We note that since $DR = R_{ox}/R_{red}$, then $R_{ox}$ can be determined from $R_{red}$ and vice-versa. In this section, we show that knowledge of $DR$ is sufficient to model how relative errors in $R$ influence $E_{GSH}$ measurement error when all parameters mapping the biosensor's ratiometric emission to $E_{GSH}$ are known except for $R_{red}$ and $R_{ox}$.

To understand the origin of the instrument-dependence of $R_{ox}$ and $R_{red}$ let us consider two imaging microscopes, M1 and M2, with identical excitation light intensity at $\lambda_1$, but with M2 having only a fraction $f$ as much excitation light intensity at $\lambda_2$ as M1. Under these conditions, a sample will exhibit the same biosensor fluorescence intensity at $\lambda_1$ in M1 and M2:

$$I_{\lambda_1,M2} = I_{\lambda_1,M1} \tag{5}$$

However, that sample will exhibit a fraction $f$ of the fluorescence intensity at $\lambda_2$ in M1 relative to M2.

$$I_{\lambda_2,M2} = f \times I_{\lambda_2,M1} \tag{6}$$

Thus, the sample's fluorescence ratio will differ across imaging microscopes by the illumination factor $f$:

$$R_{M1} = \frac{I_{\lambda_1,M1}}{I_{\lambda_2,M1}} \tag{7}$$

$$R_{M2} = \frac{I_{\lambda_1,M2}}{I_{\lambda_2,M2}} \quad (8)$$

Thus,

$$R_{M1} = f \times R_{M2} \quad (9)$$

As a result of differences in illumination across instruments, $R_{ox}$ and $R_{red}$ will be affected similarly by the illumination factor $f$. That is, under different imaging conditions:

$$R_{ox}{}' = f \times R_{ox} \quad (10)$$

$$R_{red}{}' = f \times R_{red} \quad (11)$$

and

$$R' = f \times R \quad (12)$$

In Supplementary Note 3 we showed that, in our experiments, the relative error in $R$ was independent of $R$. That is:

$$R_{observed} = R_{true} \times (1 + error) \quad (13)$$

Thus $error$ is unaffected by the illumination factor $f$:

$$R'_{observed} = f \times R_{observed} = f \times R_{true} \times (1 + error) = R_{true}{}' \times (1 + error) \quad (14)$$

Because the relative error in $R$ is equal to the relative error in $R'$, its effect on the error in $E_{GSH}$ is the same when applied to the function mapping $R$ to $E_{GSH}$ (equation 16) as it is when applied to the function mapping $R'$ to $E_{GSH}$ (equation 20), as shown below:

$$\Delta E_{GSH} = E_{observed} - E_{true} \tag{15}$$

Substituting equation (3),

$$\Delta E_{GSH} = \left[E^{\circ\prime}_{roGFP} - \frac{R_{gas}T}{2F}\ln\left(\delta_{\lambda_2}\frac{DR - \frac{R_{observed}{}'}{R_{red}{}'}}{\frac{R_{observed}{}'}{R_{red}{}'} - 1}\right)\right] - \left[E^{\circ\prime}_{roGFP} - \frac{R_{gas}T}{2F}\ln\left(\delta_{\lambda_2}\frac{DR - \frac{R_{true}{}'}{R_{red}{}'}}{\frac{R_{true}{}'}{R_{red}{}'} - 1}\right)\right] \tag{16}$$

Substituting equation (13),

$$\Delta E_{GSH} = \left[E^{\circ\prime}_{roGFP} - \frac{R_{gas}T}{2F}\ln\left(\delta_{\lambda_2}\frac{DR - \frac{R_{true}{}' \times (1 + error)}{R_{red}{}'}}{\frac{R_{true}{}' \times (1 + error)}{R_{red}{}'} - 1}\right)\right] - \left[E^{\circ\prime}_{roGFP} - \frac{R_{gas}T}{2F}\ln\left(\delta_{\lambda_2}\frac{DR - \frac{R_{true}{}'}{R_{red}{}'}}{\frac{R_{true}{}'}{R_{red}{}'} - 1}\right)\right] \tag{17}$$

Substituting with equations (11) and (14), we obtain:

$$\Delta E_{GSH} = \left[E^{\circ\prime}_{roGFP} - \frac{R_{gas}T}{2F}\ln\left(\delta_{\lambda_2}\frac{DR - \frac{f \times R_{true} \times (1 + error)}{f \times R_{red}}}{\frac{f \times R_{true} \times (1 + error)}{f \times R_{red}} - 1}\right)\right] - \left[E^{\circ\prime}_{roGFP} - \frac{R_{gas}T}{2F}\ln\left(\delta_{\lambda_2}\frac{DR - \frac{f \times R_{true}}{f \times R_{red}}}{\frac{f \times R_{true}}{f \times R_{red}} - 1}\right)\right] \tag{18}$$

$$\Delta E_{GSH} = \left[E^{\circ\prime}_{roGFP} - \frac{R_{gas}T}{2F}\ln\left(\delta_{\lambda_2}\frac{DR - \frac{R_{true} \times (1 + error)}{R_{red}}}{\frac{R_{true} \times (1 + error)}{R_{red}} - 1}\right)\right] - \left[E^{\circ\prime}_{roGFP} - \frac{R_{gas}T}{2F}\ln\left(\delta_{\lambda_2}\frac{DR - \frac{R_{true}}{R_{red}}}{\frac{R_{true}}{R_{red}} - 1}\right)\right] \tag{19}$$

Substituting equation (14),

$$\Delta E_{GSH} = \left[E^{\circ\prime}_{roGFP} - \frac{R_{gas}T}{2F}\ln\left(\delta_{\lambda_2}\frac{DR - \frac{R_{observed}}{R_{red}}}{\frac{R_{observed}}{R_{red}} - 1}\right)\right] - \left[E^{\circ\prime}_{roGFP} - \frac{R_{gas}T}{2F}\ln\left(\delta_{\lambda_2}\frac{DR - \frac{R_{true}}{R_{red}}}{\frac{R_{true}}{R_{red}} - 1}\right)\right] \tag{20}$$

$$\Delta E_{GSH} = E_{observed} - E_{true} \tag{21}$$

We conclude that knowledge of $DR$ is sufficient to model how relative errors in $R$ influence $E_{GSH}$ measurement error, when all parameters mapping the biosensor's ratiometric emission to $E_{GSH}$ are known except for $R_{red}$ and $R_{ox}$. We can therefore model how the relative errors in $R$

would influence the expected $E_{GSH}$ inaccuracy of different biosensors under otherwise identical microscopy conditions.

#### (5.2) Obtaining $\delta_{\lambda_1}$, $\delta_{\lambda_2}$, $R_{ox}$, $R_{red}$, and $DR$ parameter values for eleven roGFP-based biosensors with known midpoint potentials and fluorescence spectra

For twenty-four roGFP biosensors, we attempted to obtain the fluorescence excitation spectra of reduced and oxidized biosensor species and the biosensor $E°'$ value from published data, and from personal correspondence with authors when the spectra had not been published (Table S5-1). While $E°'$ values for all of these biosensors have been reported; the spectra of reduced and oxidized biosensor species of only eleven of these biosensors have been collected, precluding further analysis of the remaining biosensors. Spectra were digitized with WebPlotDigitizer software [10]. roGFP biosensors exhibit two peaks in their excitation spectra, centered near 410 nm (A-band) and 470 nm (B-band) (Figure S5-1, first column). Two sets of $\delta_{\lambda_1}$, $\delta_{\lambda_2}$, $R_{ox}$, $R_{red}$, and $DR$ parameter values were collected from the intensity spectra of reduced and oxidized species of each roGFP biosensor. The first set matches the bandwidths of the excitation filters in our microscopy setup, and the latter matches the wavelengths of widely used lasers. For the former, the signal at $\lambda_1$ and $\lambda_2$ were the average signals when excited over the intervals 410±15 nm and 470±10 nm, respectively. For the latter, the biosensors signal at $\lambda_1$ and $\lambda_2$ were the average signals when excited over the intervals 405±1 nm and 488±1 nm, respectively. The parameters of each biosensor were determined as described in (5.1.2). These parameters varied significantly between biosensors (Table S5-2 and Figure S5-1, second column).

#### (5.3) Predicted $E_{GSH}$ inaccuracy of eleven roGFP-based biosensors

Using the $\delta_{\lambda_2}$, $R_{ox}$, and $R_{red}$ parameters obtained in (5.2), we mapped $R$ to $E_{GSH}$ for eleven roGFP-based biosensors with known midpoint potentials and fluorescence spectra. We applied the SensorOverlord framework to determine the $E_{GSH}$ inaccuracy that we would expect to observe at each $E_{GSH}$ value with each of those biosensors, given our empirical distribution of relative errors in $R$ when measuring $E_{GSH}$ in the feeding muscles of live *C. elegans* expressing roGFP-R12 (Figure S5-1, third column).

### Table S5-1: roGFP biosensors

| Biosensor | | $E^{0'}$ (mV) | | Spectra? | |
|---|---|---|---|---|---|
| roGFP1 | [5] | -287 | [5, 11] | Yes | [5, 12] |
| roGFP2 | [5] | -272 | [5] | Yes | [1, 5, 13-16] |
| roGFP3 | [5] | -299 | [5] | Yes | [13] |
| roGFP4 | [5] | -286 | [5] | Yes | [13] |
| roGFP5 | [5] | -296 | [5] | Yes | [13] |
| roGFP6 | [5] | -280 | [5] | Yes | [13] |
| roGFP1-R1 | [7] | -269 | [7] | No | |
| roGFP1-R3 | [7] | -282 | [7] | No | |
| roGFP1-R7 | [7] | -268 | [7] | No | |
| roGFP1-R8 | [7] | -284 | [7] | No | |
| roGFP1-R9 | [7] | -278 | [7] | Yes | [17] |
| roGFP1-R10 | [7] | -284 | [7] | No | |
| roGFP1-R11 | [7] | -275 | [7] | No | |
| roGFP1-R12 | [7] | -265 | [7] | Yes | James Remington (personal communication) |
| roGFP1-R14 | [7] | -263 | [7] | No | |
| roGFP1-iL | [11] | -229 +/- 5 | [11] | Yes | [11] |
| roGFP1-iE | [11] | -236 +/- 7 | [11] | Yes | [11] |
| roGFP1-iQ | [11] | -239 +/- 6 | [11] | No | |
| roGFP1-iH | [11] | -238 +/- 4 | [11] | No | |
| roGFP1-iR | [11] | -237 +/- 5 | [11] | No | |
| roGFP1-iS | [11] | -240 +/- 3 | [11] | No | |
| roGFP1-iD | [11] | -246 +/- 1 | [11] | No | |
| roGFP2-iL | [14] | -238 | [14] | Yes | [14] |
| grx1-roGFP2 | [18] | -280 | [18] | No | |

### Table S5-2: Biosensor parameter values used for mapping $R$ to $E_{GSH}$

| Biosensor | $E^{\circ\prime}$ (mV) | Parameter set 1 $\lambda_1$ = 410 ± 15 nm and $\lambda_2$ = 470 ± 10 nm | | | | | Parameter set 2 $\lambda_1$ = 405 ± 1 nm and $\lambda_2$ = 488 ±1 nm | | | | | Spectra source |
|---|---|---|---|---|---|---|---|---|---|---|---|---|
| | | $DR$ | $\delta_{410}$ | $\delta_{470}$ | $R_{red}$ | $R_{ox}$ | $DR$ | $\delta_{405}$ | $\delta_{488}$ | $R_{red}$ | $R_{ox}$ | |
| roGFP1 | -287 | 6.08 | 1.64 | 0.27 | 3.67 | 22.33 | 3.96 | 1.66 | 0.42 | 5.25 | 20.77 | [4] |
| roGFP1 | -287 | 6.67 | 1.67 | 0.25 | 3.67 | 24.49 | 4.72 | 1.65 | 0.35 | 4.98 | 23.49 | [5] |
| roGFP1 | -287 | 4.97 | 1.24 | 0.25 | 1.42 | 7.06 | 3.99 | 1.36 | 0.34 | 2.18 | 8.69 | [12] |
| roGFP1-iE | -236 | 3.12 | 1.47 | 0.47 | 1.02 | 3.18 | 1.68 | 1.65 | 0.98 | 3.24 | 5.45 | [11] |
| roGFP1-iL | -229 | 4.13 | 1.36 | 0.33 | 2.15 | 8.87 | 2.89 | 1.39 | 0.48 | 4.09 | 11.81 | [11] |
| roGFP1-R12 * | -265 | 7.81 | 1.33 | 0.171 | 0.667 | 5.207 | | | | | | None |
| roGFP1-R12 | -265 | 5.01 | 1.30 | 0.26 | 1.42 | 7.12 | 4.04 | 1.37 | 0.34 | 2.21 | 8.93 | James Remington (personal communication) |
| roGFP1-R9 | -278 | 4.92 | 1.28 | 0.26 | 1.44 | 7.09 | 4.08 | 1.34 | 0.33 | 2.26 | 9.21 | [17] |
| roGFP2 | -272 | 6.33 | 2.22 | 0.35 | 0.15 | 0.95 | 7.89 | 3.08 | 0.39 | 0.09 | 0.71 | [5] |
| roGFP2-iL | -238 | 2.50 | 1.63 | 0.65 | 0.18 | 0.45 | 2.69 | 2.02 | 0.75 | 0.13 | 0.35 | [14] |
| roGFP3 | -299 | 4.25 | 0.98 | 0.23 | 0.97 | 4.12 | 2.95 | 1.00 | 0.34 | 1.37 | 4.04 | [13] |
| roGFP4 | -286 | 2.11 | 0.74 | 0.35 | 0.09 | 0.19 | 2.40 | 0.91 | 0.38 | 0.05 | 0.12 | [13] |
| roGFP5 | -296 | 6.92 | 1.11 | 0.16 | 1.04 | 7.20 | 4.27 | 1.15 | 0.27 | 1.87 | 7.98 | [13] |
| roGFP6 | -280 | 3.33 | 1.20 | 0.36 | 0.12 | 0.40 | 3.67 | 1.43 | 0.39 | 0.09 | 0.33 | [13] |

* $\delta_{\lambda_1}$, $\delta_{\lambda_2}$, $R_{ox}$, $R_{red}$, and $DR$ values were determined empirically in *C. elegans* expressing roGFP1-R12 in the feeding muscles [6].

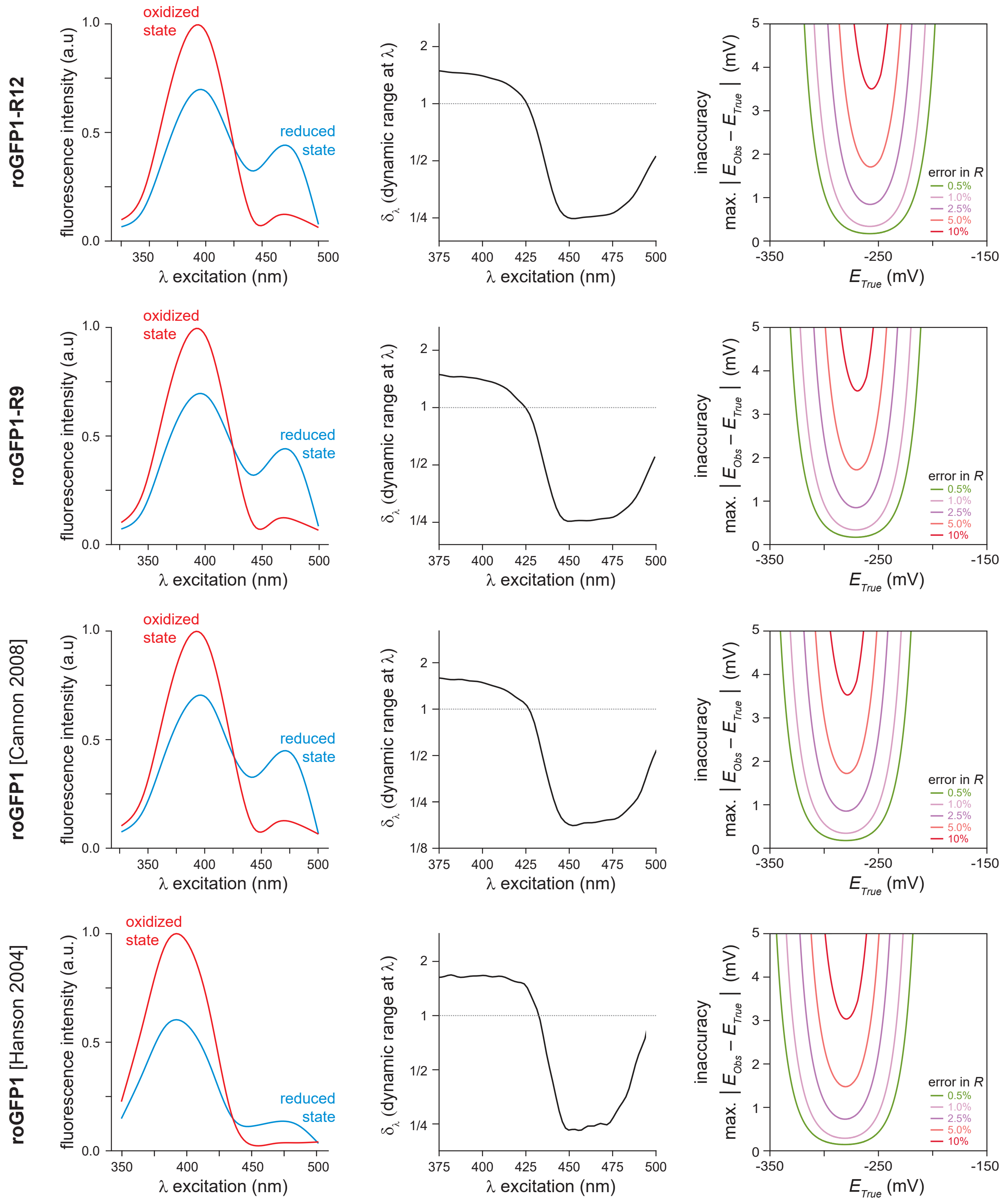


**Figure S5-1: Predicted $E_{GSH}$ measurement inaccuracy of eleven roGFP-based biosensors** In this analysis for eleven roGFP biosensors, we show in each row a plot of a roGFP biosensor's emission when excited at a wavelength $\lambda$ in the oxidized and reduced states (column 1); a plot of the ratio of those emissions, the biosensor's dynamic range $\delta_\lambda$, at each wavelength $\lambda$ (column 2); and a plot of the predicted $E_{GSH}$ measurement inaccuracy with that biosensor at each wavelength $\lambda$ for different relative error in $R$ values (column 3).

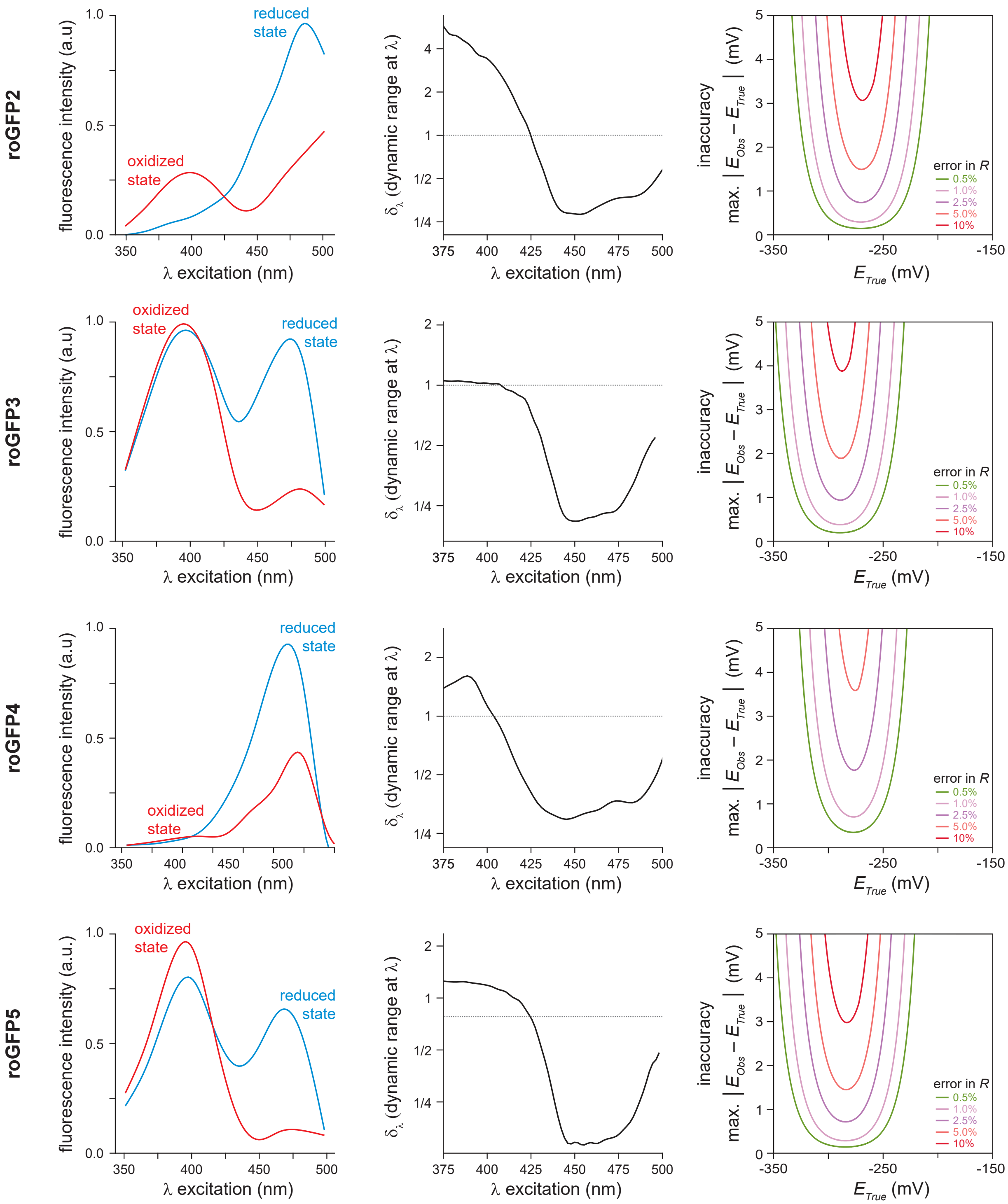
roGFP2
roGFP3
roGFP4
roGFP5
fluorescence intensity (a.u)
fluorescence intensity (a.u.)
λ excitation (nm)
oxidized state
reduced state
$\delta_\lambda$ (dynamic range at λ)
inaccuracy
max. | $E_{Obs} - E_{True}$ | (mV)
$E_{True}$ (mV)
error in $R$
0.5%
1.0%
2.5%
5.0%
10%

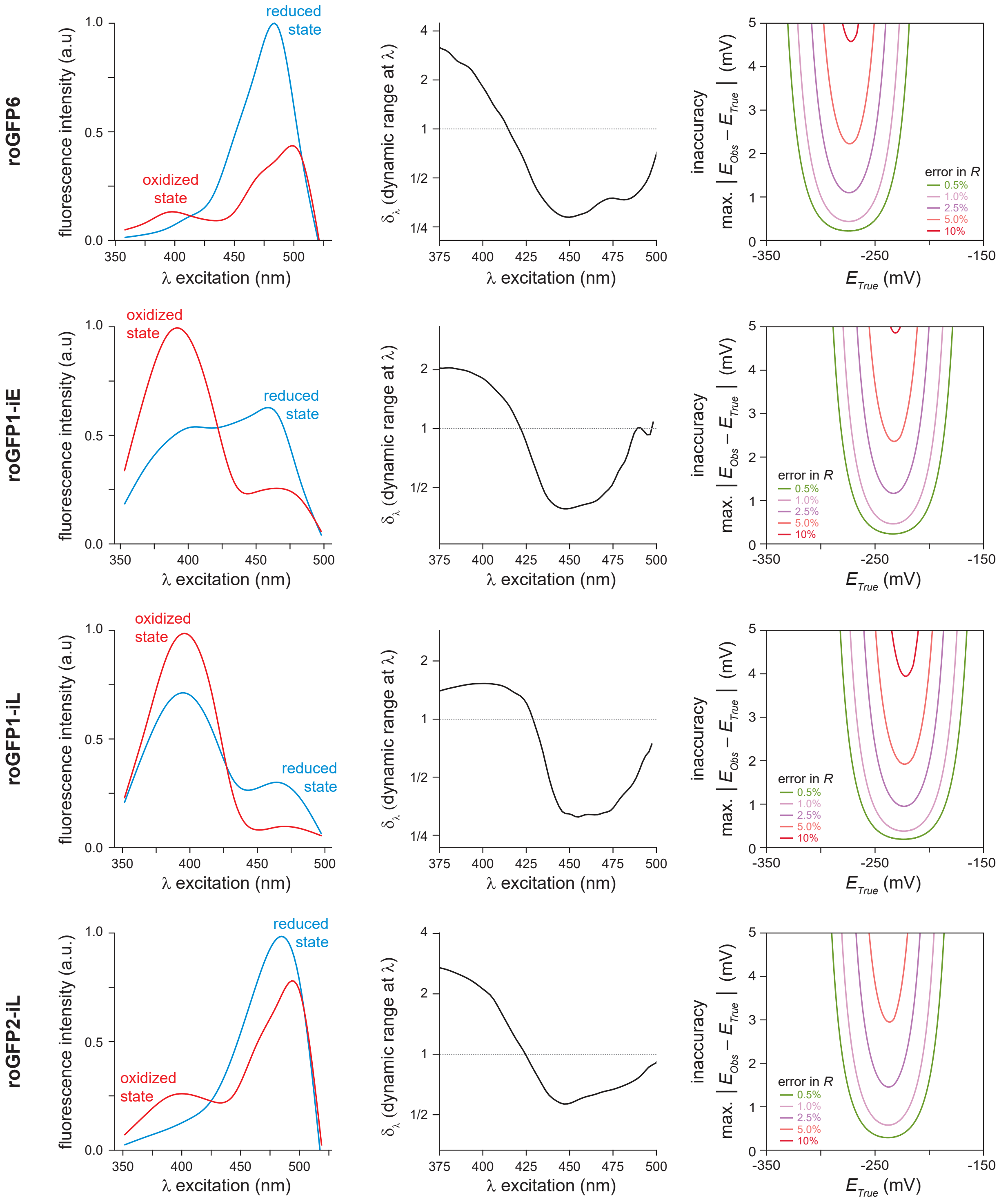

roGFP6
roGFP1-iE
roGFP1-iL
roGFP2-iL
fluorescence intensity (a.u)
fluorescence intensity (a.u.)
λ excitation (nm)
reduced state
oxidized state
δλ (dynamic range at λ)
inaccuracy
max. | E_Obs − E_True | (mV)
E_True (mV)
error in R
0.5%
1.0%
2.5%
5.0%
10%

**Supplementary Note 6**

In this note, we compared biochemical and biophysical properties of roGFP biosensors that are important predictors of biosensor $E_{GSH}$ accuracy.

**(6.1) Comparing the overall dynamic range and effective midpoint potential of roGFP biosensors**

In Supplementary Note 1.3 we derived the function mapping the ratiometric emission ($R$) of roGFP biosensors to the reduction potential of the glutathione-glutathione disulfide couple $E_{GSH}$:

$$E_{GSH} = E_{roGFP} = E^{\circ\prime}_{roGFP} - \frac{R_{gas}T}{2F} \ln \left(\delta_{\lambda_2} \frac{R_{ox} - R}{R - R_{red}}\right) \quad (1)$$

In Supplementary Note 4.2 we modeled the sensitivity of $E_{GSH}$ measurement inaccuracy to changes in the values of the parameters in this map that quantify the biochemical and biophysical properties of the biosensor and instrumentation. Changes in two of these parameters shift $E_{GSH}$ by constant amounts: the biosensor's standard midpoint potential $E^{\circ\prime}_{roGFP}$ and the biosensor's dynamic range in the second excitation wavelength $\delta_{\lambda_2}$. We can aggregate these two effects into a single effective midpoint potential $E^{\circ\prime}_{effective}$:

$$E^{\circ\prime}_{effective} = E^{\circ\prime}_{roGFP} - \frac{R_{gas}T}{2F} \ln \delta_{\lambda_2} \quad (2)$$

$$E_{GSH} = E^{\circ\prime}_{effective} - \frac{R_{gas}T}{2F} \ln \frac{R_{ox} - R}{R - R_{red}} \quad (3)$$

In Supplementary Note 4.2, we also found that the optimal strategy for minimizing $E_{GSH}$ inaccuracy with a given biosensor and error level in $R$ is to select excitation wavelengths that maximize the biosensor's dynamic range $DR$.

Both $E^{\circ\prime}_{effective}$ and $DR$ depend on the choice of excitation wavelengths $\lambda_1$and $\lambda_2$. As a result, the values of these parameters can vary across experimental settings. In Supplementary Note 5.2 we derived the values of $DR$ and $\delta_{\lambda_2}$ for each of the eleven roGFP biosensors with known fluorescence spectra. These values were determined for two illumination settings: one matched the bandwidths of the excitation filters in our microscopy set up and the other matched the wavelengths of widely used lasers. Using each of those two sets of parameters, we inspected how $E^{\circ\prime}_{effective}$ and $DR$ covary across roGFP biosensors (Figure S6-1). This analysis suggested that no roGFP biosensor had $DR$ values above 5 and $E^{\circ\prime}_{effective}$ values above -275 mV under either imaging modality (Figure S6-1a,b). Most biosensors were predicted to have higher $DR$ values using parameters for 410±15 nm and 470±10 nm illumination (Figure S6-1a) than using parameters for 405 nm and 488 nm illumination (Figure S6-1b).

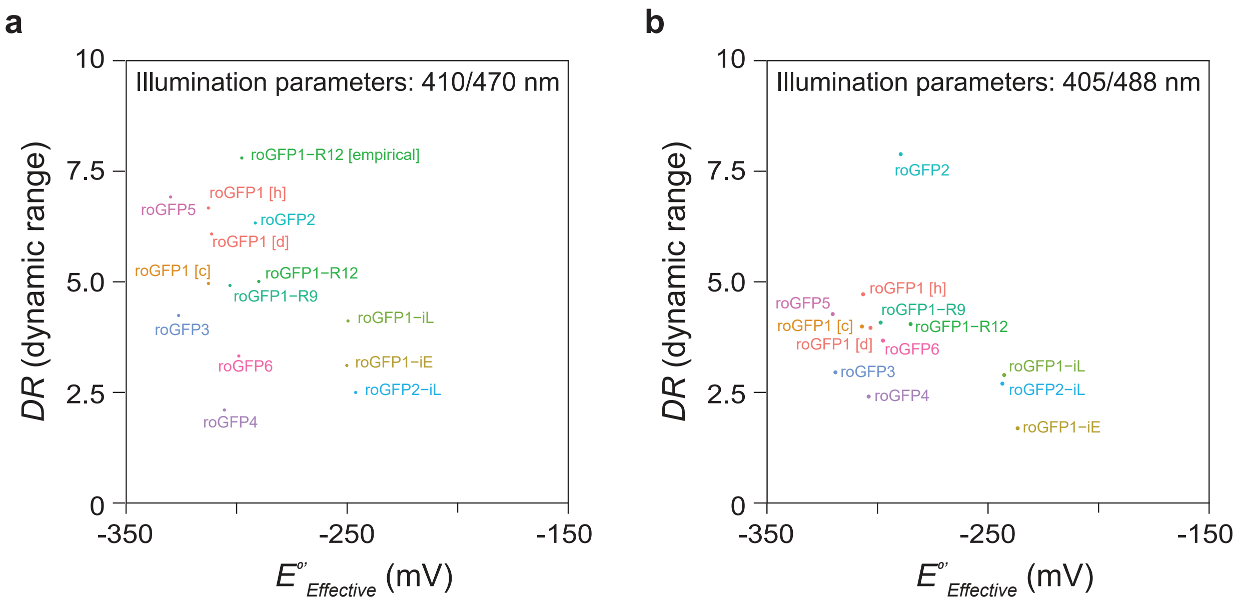


**Figure S6-1: $E^{\circ\prime}_{effective}$ and $DR$ covary across roGFP biosensors**

The values of $E^{\circ\prime}_{effective}$ and $DR$ depend on the choice of excitation wavelengths $\lambda_1$and $\lambda_2$ and therefore can vary across experimental settings. The scatterplots in panels **a** and **b**, show the covariation of $E^{\circ\prime}_{effective}$ and $DR$ of the eleven roGFP biosensors with known fluorescence spectra for two experimental settings. In panel **a**, these values were determined for the illumination setting matching the bandwidths of the excitation filters in our microscopy set up; while, in panel **b**, these values were determined for the illumination setting matching the wavelengths of widely used 405 nm and 488 nm lasers.

**Supplementary Note 7**

In this note, we build on the framework described in Supplementary Note 1 to derive (7.1) the map from fluorescence-ratio ($R$) measurements to pH in two-state ratiometric pH biosensors; and (7.2) the more general map from $R$ to ligand concentration in ligand-binding two-state ratiometric biosensors.

**(7.1) Mapping *R* to pH in two-state ratiometric pH biosensors**

(7.1.1) General chemistry of pH-sensitive two-state ratiometric biosensors

The chromophore of GFP and the chromophores of nearly all fluorescent proteins are formed via a spontaneous series of reactions within an internal X-Tyr-Gly sequence [19, 20]. The phenolic hydroxyl group in the chromophore in each of these proteins, derived from the Tyr side chains, is a weak acid. Protonated and deprotonated chromophore forms exhibit different fluorescence-excitation peaks. For example, in the case of GFP, the protonated chromophore exhibits an excitation peak at 395 nm while the deprotonated form exhibits a peak at 475 nm [2]. The protonation state of the chromophore of many fluorescent proteins, including GFP and mRFP, is relatively insensitive to the pH of the surrounding environment [21]. In contrast, many mutant variants derived from GFP and other fluorescent proteins have chromophores whose protonation state is sensitive to the pH of the surrounding environment and can, therefore, be used as pH biosensors [2].

For many pH biosensors, the protonated chromophore state exhibits very poor fluorescence emission. Therefore, the fluorescence emission of an ensemble of such pH biosensors depends on the number of deprotonated biosensors and the fraction of those biosensors in the deprotonated state. Because both of these quantities are generally unknown *in vivo*, these intensity-based biosensors can be useful reporters of changes in pH but are not useful for absolute pH measurement [20].

Ratiometric pH-sensitive biosensors overcome this limitation, enabling calculation of the fraction of deprotonated biosensors from the ratio of the biosensor's fluorescence excitation and/or emission at two different wavelengths ($R$), as we describe in Supplementary Note 1.2.

Knowledge of the fraction of deprotonated biosensors is a first step towards measuring pH with pH-sensitive biosensors, which also requires knowledge of the biosensor's tendency to become protonated and deprotonated. The balance between protonated and deprotonated chromophore forms of a fluorescent biosensor is described by the weak acid equation:

$$[Biosensor^{-}] + [H^{+}] \rightleftarrows [BiosensorH] \tag{1}$$

where $BiosensorH$ and $Biosensor^{-}$ are the biosensors with protonated and deprotonated chromophores, respectively. At equilibrium, the balance among the concentrations of these species is described by the biosensor's chromophore acid dissociation constant $K_a$:

$$K_a = \frac{[Biosensor^{-}] \times [H^{+}]}{[BiosensorH]} \tag{2}$$

$K_a$is determined by the local environment surrounding the chromophore and is therefore a property of each biosensor. Solving for $[H^{+}]$, we obtain the Henderson-Hasselbach equation:

$$pH = pK_a + log_{10} \frac{[Biosensor^{-}]}{[BiosensorH]} \tag{3}$$

In the section (7.1.3) we use this equation to calculate pH given knowledge of both the biosensor's $K_a$and the fraction of deprotonated biosensors.

<u>(7.1.2) Deriving the map between a biosensor's fluorescence ratio and the fraction of deprotonated biosensors</u>

The derivation of the map between $R$ and $P(Biosensor^{-})$ builds on the framework developed for two-state ratiometric biosensors in Supplementary Note 1.2. For roGFPs, the A and B biosensor states refer to the oxidized and reduced biosensor species. Here, these two states refer to the deprotonated and protonated biosensor species. Starting with equation 12 from Supplementary Note 1.2, and substituting $Biosensor^{-}$ and $BiosensorH$ for the biosensor's A and B states, we obtain the map between $R$ and $P(Biosensor^{-})$:

$$P(Biosensor^{-}) = \frac{R - R_{BiosensorH}}{R - R_{BiosensorH} + \delta_{\lambda_2}(R_{Biosensor^{-}} - R)} \tag{4}$$

<u>(7.1.3) Deriving the map between a biosensor's fluorescence ratio and pH</u>

The concentrations of protonated and deprotonated biosensor species are constrained by mass balance:

$$[Biosensor] = [Biosensor^{-}] + [BiosensorH] \quad (5)$$

Substituting the Henderson-Hasselbach equation we obtain:

$$pH = pK_a - log_{10}\frac{1 - P(Biosensor^{-})}{P(Biosensor^{-})} \quad (6)$$

Substituting the map between $R$ and $P(Biosensor^{-})$ we obtain:

$$pH = pK_a - log_{10}(\delta_{\lambda_2}\frac{R_{Biosensor^{-}} - R}{R - R_{BiosensorH}}) \quad (7)$$

This is the map between a biosensor's fluorescence ratio and the pH of the surrounding environment under equilibrium conditions.

We note that this map has the same general form as that from $R$ to $E_{GSH}$ in roGFP biosensors (see Supplementary Note 1.3). In the next section, we extend this framework to ligand-binding biosensors.

**(7.2) Mapping *R* to ligand concentration in two-state ratiometric ligand-binding biosensors**

7.2.1. General chemistry of ligand-binding biosensors

The protonation-deprotonation reaction of a pH-sensitive biosensor:

$$[Biosensor^{-}] + [H^{+}] \rightleftarrows [BiosensorH] \quad (8)$$

is a special case of the more general binding-unbinding reaction of a single ligand and a single biosensor:

$$[Biosensor] + [Ligand] \rightleftarrows [Biosensor - Ligand] \quad (9)$$

where $[Biosensor]$, $[Ligand]$, and $[Biosensor - Ligand]$ represent the concentrations of unbound free biosensor, unbound free ligand, and biosensor-ligand complex.

At equilibrium, the balance among the concentrations of $Biosensor$, $Ligand$, and $Biosensor - Ligand$ species is described by the dissociation constant $K_d$:

$$K_d = \frac{[Biosensor] \times [Ligand]}{[Biosensor - Ligand]} \tag{10}$$

In section 7.2.2, we derive the map from a biosensor's fluorescence ratio (*R*) to the concentration of its ligand for two-state ratiometric ligand-binding biosensors.

<u>(7.2.2) Deriving the map between a biosensor's fluorescence ratio and ligand concentration</u>

Specific two-state ratiometric biosensors exist for many ligands, including Histidine, $NAD^+$, NADH, and NADPH. The derivation of the map between $R$ and $[Ligand]$ mirrors that between $R$ and pH in two-state ratiometric pH biosensors. Starting with equation 12 from Supplementary Note 1.2, and substituting $Biosensor$ and $Biosensor - Ligand$ for the biosensor's A and B states, we obtain the map between $R$ and $P(Biosensor)$:

$$P(Biosensor) = \frac{R - R_{Biosensor-Ligand}}{R - R_{Biosensor-Ligand} + \delta_{\lambda_2}(R_{Biosensor} - R)} \tag{11}$$

Solving for $[Ligand]$ we obtain:

$$[Ligand] = 10^{-pK_d + log_{10}\frac{1-P(Biosensor)}{P(Biosensor)}} \tag{12}$$

Substituting the map between $R$ and $P(Biosensor)$ we obtain:

$$[Ligand] = 10^{-pK_d + log_{10}(\delta_{\lambda_2}\frac{R_{Biosensor}-R}{R-R_{Biosensor-Ligand}})} \tag{13}$$

This is the map between a biosensor's fluorescence ratio and the free ligand concentration of the surrounding environment under equilibrium conditions.

**Supplementary Note 8**

In Supplementary Note 7, we derived the functions mapping the ratiometric emission ($R$) of two-state biosensors to the pH in the environment surrounding those biosensors under equilibrium conditions. Because those maps have the same general form as those from $R$ to $E_{GSH}$ in roGFP biosensors (see Supplementary Note 1.3), they also have a similar dependence on biochemical and biophysical parameters quantifying the properties of the biosensors and the instrumentation used to measure their fluorescence (see Supplementary Note 5.1). Knowledge of those parameters enabled us to model how the precision of $R$ measurements with a biosensor determines the range of $E_{GSH}$ values that is possible to measure with that biosensor at a specific inaccuracy level (see Supplementary Note 5.2 and 5.3). In this note, we build on that scheme to model how the precision of $R$ measurements with a biosensor determines the range of pH values that is possible to measure with that biosensor at a given inaccuracy level. We (8.1) estimate the values of the minimal set of parameters and constraints between parameters needed to predict the influence of relative errors in $R$ on nine pH biosensors with known $pK_a$ and fluorescence spectra; and (8.2) determine the pH inaccuracy we would expect to observe with each of those biosensors if we selected optimal excitation or emission filters for each biosensor and measured $R$ at the same precision as our $R$ measurements in the feeding muscles of live *C. elegans* with roGFP-R12. In Supplementary Note 9 we extend this analysis to two-state ligand-binding ratiometric biosensors.

**(8.1) Obtaining the values of the parameters that map $R$ to pH for nine two-state ratiometric pH biosensors**

(8.1.1) Overall approach

In Supplementary Note 7.1.3 we derived the function mapping a biosensor's fluorescence ratio ($R$) and the pH of the environment surrounding the biosensor under equilibrium conditions.

$$pH = pK_a - log_{10}(\delta_{\lambda_2}\frac{R_{Biosensor^-} - R}{R - R_{BiosensorH}}) \quad (1)$$

This map depends on biochemical and biophysical parameters. The biochemical parameter is the biosensor's chromophore acid dissociation constant $K_a$. This parameter is often known from published reports; however, in many cases the values reported do not represent true $K_a$ values as they are extracted from pH-titrations of the biosensor's fluorescence ratio, which do not necessarily have a linear relationship with the fraction of protonated biosensors (see Supplementary

Note 2). Therefore, we show how these reported effective $K_a$ values can be corrected with knowledge of some of the biosensor's biophysical parameters (section 8.1.2).

The biophysical parameters required to map $R$ to pH include the biosensor's dynamic range in the second wavelength $\delta_{\lambda_2}$, the ratiometric emission of an ensemble of protonated biosensors $R_{BiosensorH}$, and the ratiometric emission of an ensemble of deprotonated biosensors $R_{Biosensor^-}$. These three parameters can be determined empirically, but in most cases they are not known. The value of $\delta_{\lambda_2}$ and the biosensor's overall dynamic range $DR$ (equal to the $R_{Biosensor^-}/R_{BiosensorH}$ ratio) can be determined from the fluorescence spectra of protonated and deprotonated biosensors, similar to the case of reduced and oxidized biosensor species (see Supplementary Note 5.1.2). In the case of pH biosensors, however, published reports often underestimate the biosensor's dynamic range because these reports only include the spectra of biosensor species that are not fully protonated or fully deprotonated, often because the biosensors begin to unfold at pH > 10 [3]. We show how these published spectra and knowledge of the biosensor's $K_a$ value can be used to estimate the spectra of protonated and deprotonated biosensor species, leading to more accurate estimates of the biosensor's dynamic range at each wavelength (section 8.1.3).

We then estimate the values of the minimal set of parameters and constraints between parameters needed to predict the influence of relative errors in $R$ on nine pH biosensors with known $pK_a$ and fluorescence spectra (section 8.1.4). As discussed in Supplementary Note 5.1.3 for the map from $R$ to $E_{GSH}$, mapping the $R$ values obtained with a given instrument to pH requires knowledge of the values of both $R_{BiosensorH}$ and $R_{Biosensor^-}$ in that instrument; however, when all other parameters are known, knowledge of the $R_{Biosensor^-}/R_{BiosensorH}$ ratio is sufficient for modeling how relative errors in $R$ would influence the expected pH inaccuracy of different biosensors under otherwise identical microscopy conditions.

<u>(8.1.2) Obtaining the $pKa$ of a biosensor from a pH-titration of its fluorescence ratio</u>

The balance among the concentrations of protonated and deprotonated species of a pH biosensor is quantified by the biosensor's chromophore acid dissociation constant $K_a$. Empirically, $K_a$ is often determined as the point in a pH titration where deprotonated and protonated biosensor species have equal concentrations. However, studies often report the pH value where the biosensor's fluorescence ratio is half-way between the values corresponding to protonated and deprotonated biosensors. These "effective $pK_a$" ($pK_{effective}$) values do not represent true $pK_a$ values as they are extracted from pH-titrations of the biosensor's fluorescence ratio, and not of the concentration of specific biosensor species. To convert into $pK_a$ values the reported $pK_{effective}$ values, we substitute $R$ = $(R_{Biosensor^-} - R_{BiosensorH})/2$ in equation 1:

$$pH = pK_{effective} = pK_a - log_{10}\delta_{\lambda_2} \quad (2)$$

where $\delta_{\lambda_2}$ is the biosensor's dynamic range in the second wavelength used in the titration. Therefore:

$$pK_a = pK_{effective} + log_{10}\delta_{\lambda_2} \quad (3)$$

(8.1.3) Obtaining $\delta_{\lambda_2}$ and $DR$ from the excitation spectra of pH biosensor species that are not fully protonated or deprotonated

Published reports often only include the spectra of biosensor species that are not fully protonated or fully deprotonated and therefore underestimate the biosensor's dynamic range. Here, we show how to estimate the biosensor's dynamic range using published spectra obtained at two pH values and knowledge of the biosensor's $pK_a$ value.

In Supplementary Note 1.2.1, we showed that in a mixed population of biosensors the total observed fluorescence intensity will be the weighted average of the intensities of populations of biosensors in each state:

$$I_\lambda = P(A)\, I_{\lambda,A} + [1 - P(A)]\, I_{\lambda,B} \quad (4)$$

Where $\lambda$ represents an excitation- and emission-wavelength pair, A and B are the two biosensor states, $P(A)$ is the fraction of biosensors in state A, and $I_{\lambda,A}$ and $I_{\lambda,B}$ are, respectively, the total observed intensities of ensembles of biosensors in states A and B. For pH biosensors, the A state corresponds to the deprotonated state and the B state corresponds to the protonated state.

To obtain $I_{\lambda,Biosensor^-}$ and $I_{\lambda,BiosensorH}$ from $I_\lambda$ measurements at two different pH values, x and y, we first calculate $P(Biosensor^-)$ at those pH values:

$$a = P(Biosensor^{-}, pH = x) = \frac{1}{1 + 10^{(pK_a - x)}} \tag{5}$$

$$b = P(Biosensor^{-}, pH = y) = \frac{1}{1 + 10^{(pK_a - y)}} \tag{6}$$

Substituting equations 5 and 6 in equation 4, we obtain:

$$I_{\lambda,x} = \ a\, I_{\lambda,Biosensor^{-}} + (1 - a)\, I_{\lambda,BiosensorH} \tag{7}$$

$$I_{\lambda,y} = \ b\, I_{\lambda,Biosensor^{-}} + (1 - b)\, I_{\lambda,BiosensorH} \tag{8}$$

Therefore:

$$\frac{I_{\lambda,x} - a\, I_{\lambda,Biosensor^{-}}}{1 - a} = \frac{I_{\lambda,y} - b\, I_{\lambda,Biosensor^{-}}}{1 - b} \tag{9}$$

$$\frac{I_{\lambda,x} - (1 - a)\, I_{\lambda,BiosensorH}}{a} = \frac{I_{\lambda,y} - (1 - b)\, I_{\lambda,BiosensorH}}{b} \tag{10}$$

Solving for $I_{\lambda,Biosensor^{-}}$ and $I_{\lambda,BiosensorH}$:

$$I_{\lambda,Biosensor^{-}} = \frac{(1 - b)\, I_{\lambda,x} - (1 - a)\, I_{\lambda,y}}{a - b} \tag{11}$$

$$I_{\lambda,BiosensorH} = \frac{b\, I_{\lambda,x} - a\, I_{\lambda,y}}{b - a} \tag{12}$$

(8.1.4) $pK_a$, $\delta_{\lambda_1}$, $\delta_{\lambda_2}$, $R_{BiosensorH}$ and $R_{Biosensor^-}$, and $DR$ parameter values for nine two-state ratiometric pH biosensors

For fifteen ratiometric two-state pH biosensors, we attempted to obtain the biosensor's $pK_a$ value and the fluorescence excitation spectra of protonated and deprotonated biosensor species from published data (Table S8-1). Of the five biosensors where the $pK_a$ was unknown, the $pK_{effective}$ was known for two, Sypher and Sypher3s. For these biosensors, we obtained an estimate of their $pK_a$ from pH-titrations of their fluorescence ratios, following the scheme described in section 8.1.2 (Table S8-2). The fluorescence excitation spectra of protonated and deprotonated biosensor species have not been reported for four of the fifteen pH biosensors (Table S8-1).

For the nine biosensors with available spectra and known or estimated $pK_a$, spectra collected at different pH values were digitized with the WebPlotDigitizer software [10], and the spectra of fully protonated and fully deprotonated biosensor states were estimated following the scheme described in section 8.1.3. In nine cases these spectra differed only slightly from the empirical measurements (Figures S8-1,2,3), because these measurements had been obtained at pH values at least one pH unit above and one pH unit below the $pK_a$ of the biosensor. The exception was SypHer, a biosensor where all spectra were collected at pH values below the $pK_a$. The estimate of this biosensor's $\delta_{\lambda_2}$and $DR$ were therefore considered of insufficient quality and the biosensor was not studied further.

For each of the eight remaining pH biosensors, $\delta_{\lambda_1}$, $\delta_{\lambda_2}$, $R_{BiosensorH}$, $R_{Biosensor^-}$, and $DR$ parameter values were collected from the estimated fluorescence intensity spectra of fully protonated and fully deprotonated biosensor states (Table S8-2). For each biosensor, we chose excitation-emission-wavelength pairs, $\lambda_1$ and $\lambda_2$, seeking to maximize $DR$ and to match the wavelengths at which the biosensor's fluorescence intensity peaked. We used fluorescence intensity signal averaged over 20 nm intervals centered around $\lambda_1$ and $\lambda_2$ to calculate $\delta_{\lambda_1}$, $\delta_{\lambda_2}$, $R_{BiosensorH}$, $R_{Biosensor^-}$, and $DR$. The values of these parameters for each biosensor were determined as described in (8.1). These parameters varied significantly between biosensors (Table S8-2 and Figures S8-1,2,3 second column).

#### (8.2) Predicted pH inaccuracy of eight ratiometric two-state pH biosensors

Using the $pK_a$, $\delta_{\lambda_2}$, $R_{BiosensorH}$, and $R_{Biosensor^-}$ parameters obtained in (8.1), we mapped $R$ to pH for eight ratiometric two-state pH biosensors. We applied the SensorOverlord framework to determine the pH inaccuracy that we would expect to observe at each pH value with each biosensor, given our empirical distribution of relative errors in $R$ when measuring $E_{GSH}$ in the feeding muscles of live *C. elegans* expressing roGFP-R12. This analysis was applied to dual-excitation red-fluorescent pH biosensors (Figure S8-1, third column), dual-excitation green-fluorescent pH biosensors (Figure S8-2, third column), and single-excitation dual-emission biosensors (Figure S8-3, third column). The $E^2$GFP biosensor can be used in two different modalities, dual-excitation green-fluorescence and single-excitation dual-emission. The differences in the biosensor's $\delta_{\lambda_2}$ and $DR$ parameter values in each imaging modality lead to differences in the predicted pH inaccuracy of this biosensor under each imaging modality.

### Table S8-1: two-state ratiometric pH biosensors

| Biosensor | | $pK_a$ | $pK_{effective}$ | | Spectra? | |
|---|---|---|---|---|---|---|
| deGFP1 | [3] | 8.02 | | [3] | Yes | [3, 13] |
| deGFP2 | [3] | 7.25 | | [3] | No | |
| deGFP3 | [3] | 6.86 | | [3] | No | |
| deGFP4 | [3] | 7.37 | | [3] | Yes | [3, 13] |
| ClopHensor | [22] | 6.78 | | [22] | Yes | [22] |
| $E^1$GFP | [23] | No | | | No | |
| $E^2$GFP | [24] | 7.01<br>6.78 | | [23, 24] | Yes | [24] |
| ratiometric-pHluorin | [25] | No | | | Yes | [25] |
| pHluorin2 | [26] | No | | | Yes | [26] |
| mCherryEA | [27] | 7.8 | | [27] | Yes | [27] |
| pHRed | [28] | 7.8 | | [28] | Yes | [28] |
| mKeima | [28] | 7.8 | | [28] | Yes | [28] |
| SypHer3s | [29] | No | 7.8 | [29] | Yes | [29] |
| SypHer-2 | [30] | 8.1 | | [30] | No | |
| SypHer | [31] | No | 8.71 | [31] | Yes | [31] |

### Table S8-2: Biosensor parameter values used for mapping $R$ to pH

| Biosensor | Imaging modality | $\lambda_1 \pm 10\ nm$ | $\lambda_2 \pm 10\ nm$ | $pK_a$ | $DR$ | $\delta_{\lambda_1}$ | $\delta_{\lambda_2}$ | $R_{BiosensorH}$ | $R_{Biosensor^-}$ |
|---|---|---|---|---|---|---|---|---|---|
| mCherryEA | dual excitation | 455 | 585 | 7.8 | 9.87 | 3.16 | 0.32 | 0.30 | 2.96 |
| pHRed | dual excitation | 440 | 585 | 7.8 | 62.6 | 6.26 | 0.10 | 0.15 | 9.39 |
| mKeima | dual excitation | 440 | 560 | 7.8 | 21.3 | 2.13 | 0.10 | 0.95 | 20.2 |
| $E^2$GFP | dual excitation | 480 | 440 | 6.78 | 15.7 | 1.26 | 0.08 | 0.32 | 5.02 |
| ClopHensor | dual excitation | 488 | 430 | 6.78 | 8.64 | 2.85 | 0.33 | 1.16 | 10.0 |
| SypHer3s | dual excitation | 495 | 410 | 8.5 * | 414 | 53.9 | 0.13 | 0.19 | 78.7 |
| $E^2$GFP | dual emission | 525 | 505 | 6.78 | 5.02 | 4.17 | 0.83 | 0.82 | 4.12 |
| deGFP4 | dual emission | 515 | 460 | 7.37 | 62.5 | 10.0 | 0.16 | 0.47 | 29.4 |
| deGFP1 | dual emission | 515 | 460 | 8.02 | 47.3 | 10.9 | 0.23 | 0.58 | 27.4 |

* $pK_a$ derived from $pK_{effective}$ as described in section 8.1.4.

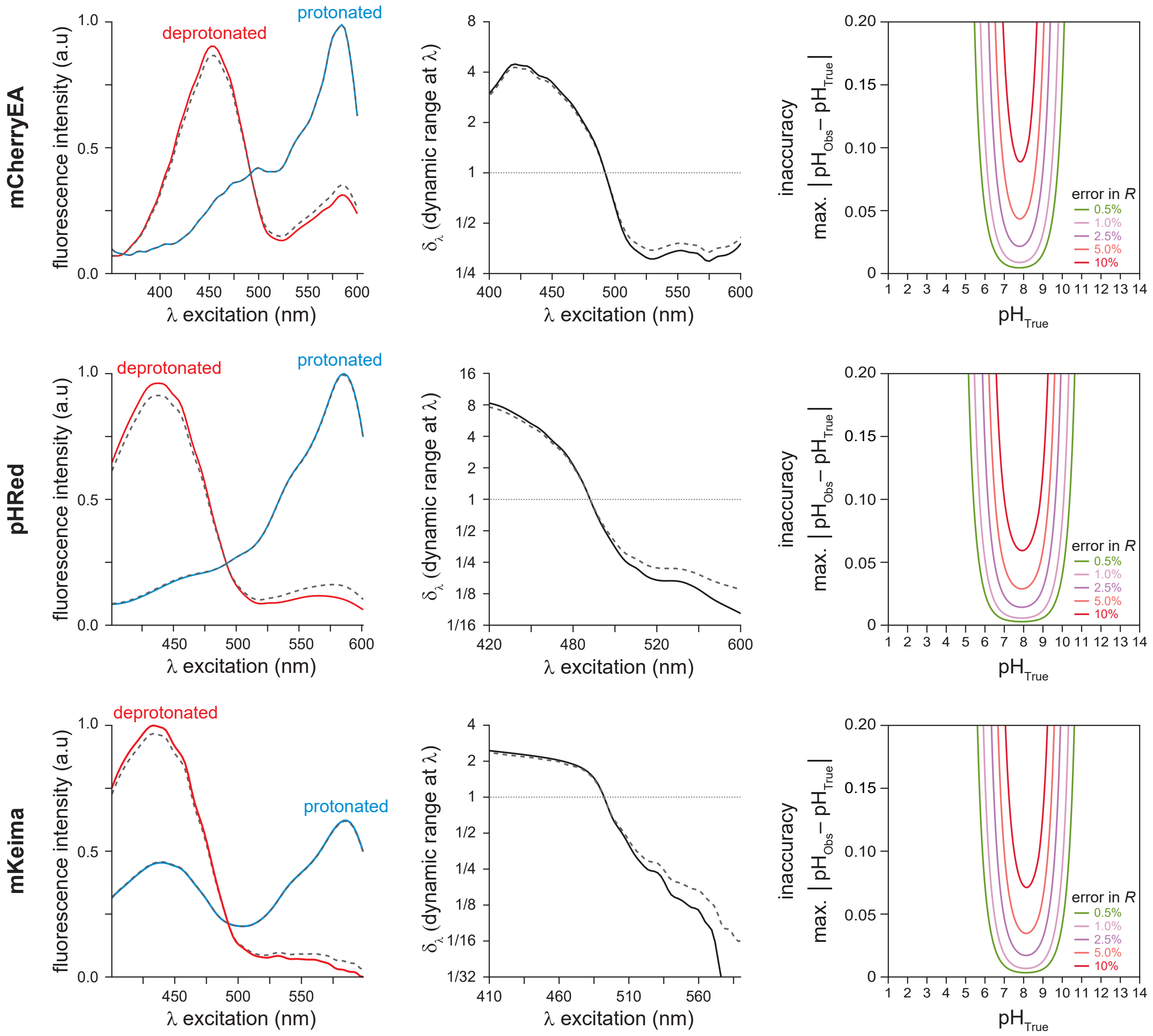


**Figure S8-1: Predicted pH inaccuracy of dual-excitation red-fluorescent pH biosensors**

In this analysis of three dual-excitation red-fluorescent pH biosensors, we show in each row a plot of the biosensor's predicted emission when excited at a wavelength $\lambda$ in the fully deprotonated and protonated states (solid lines, column 1) and the observed emissions of mostly deprotonated and mostly protonated biosensor mixtures (dotted lines, column 1); a plot of the biosensor's dynamic range $\delta_\lambda$, the predicted ratio of the emissions of the fully deprotonated and protonated biosensor states (solid line, column 2) and the apparent $\delta_\lambda$ derived from the ratio of the observed emissions of mostly deprotonated and mostly protonated biosensor mixtures (dotted line, column 2); and a plot of the predicted pH measurement inaccuracy with that biosensor at each wavelength $\lambda$ for different relative errors in $R$ values (column 3).

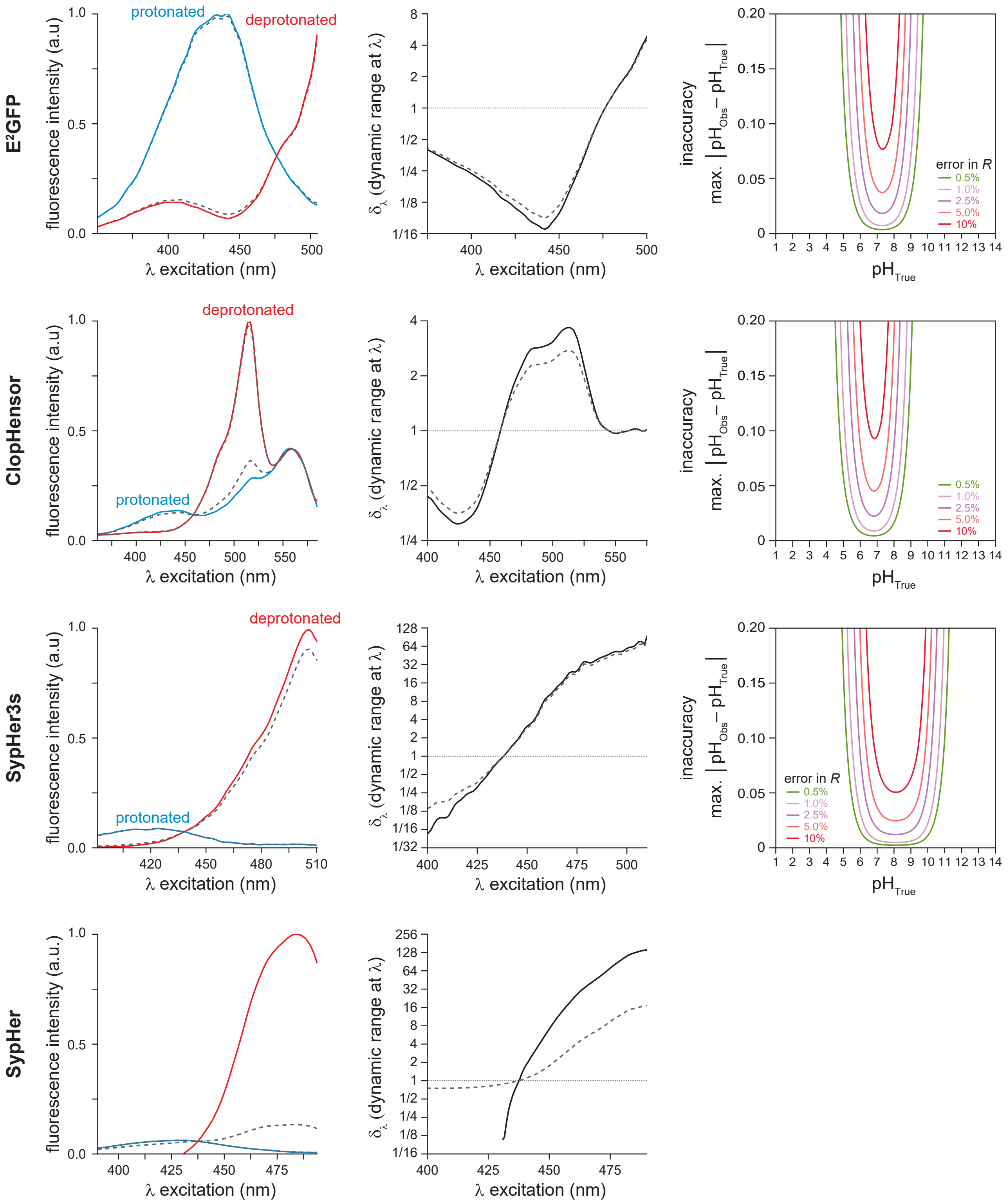


**Figure S8-2: Predicted pH inaccuracy of dual-excitation green-fluorescent pH biosensors**

In this analysis of four dual-excitation green-fluorescent pH biosensors, we show in each row a plot of the biosensor's predicted emission when excited at a wavelength $\lambda$ in fully deprotonated and protonated states (solid lines, column 1) and the observed emissions mostly deprotonated and mostly protonated biosensor mixtures (dotted lines, column 1); a plot of the biosensor's dynamic range $\delta_\lambda$, the predicted ratio of the emissions of fully deprotonated and protonated biosensor states (solid line, column 2) and the apparent $\delta_\lambda$ derived from the ratio of the observed emissions of mostly deprotonated and mostly protonated biosensor mixtures (dotted line, column 2); and a plot of the predicted pH measurement inaccuracy with that biosensor at each wavelength $\lambda$ for different relative errors in $R$ values (column 3).

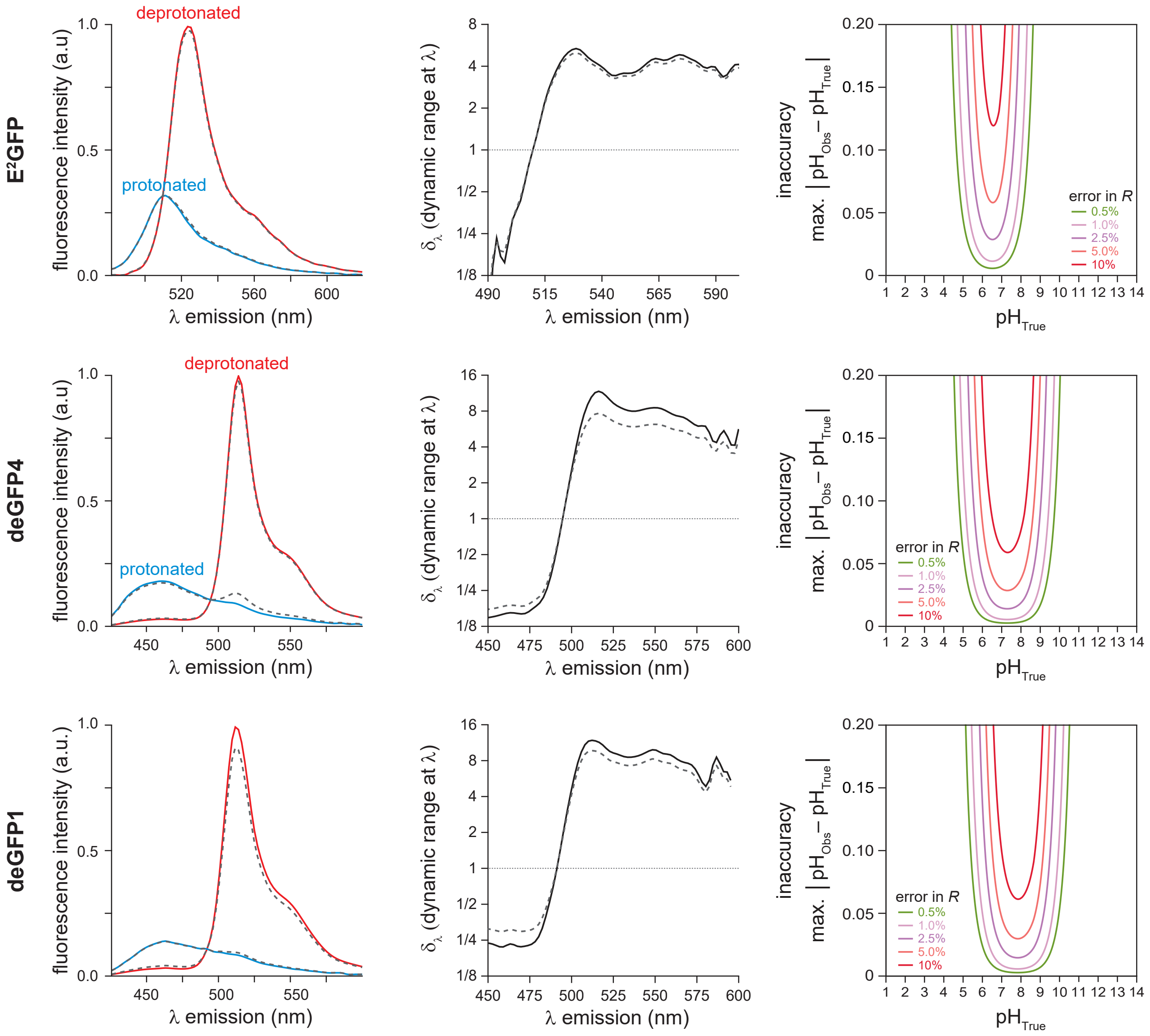


**Figure S8-3: Predicted pH inaccuracy of single-excitation dual-emission pH biosensors**

In this analysis of three single-excitation dual-emission pH biosensors, we show in each row a plot of the biosensor's predicted emission when excited at a wavelength $\lambda$ in fully deprotonated and protonated states (solid lines, column 1) and the observed emissions mostly deprotonated and mostly protonated biosensor mixtures (dotted lines, column 1); a plot of the biosensor's dynamic range $\delta_\lambda$, the predicted ratio of the emissions of fully deprotonated and protonated biosensor states (solid line, column 2) and the apparent $\delta_\lambda$ derived from the ratio of the observed emissions of mostly deprotonated and mostly protonated biosensor mixtures (dotted line, column 2); and a plot of the predicted pH measurement inaccuracy with that biosensor at each wavelength $\lambda$ for different relative errors in $R$ values (column 3).

**Supplementary Note 9**

In Supplementary Note 7, we derived the functions mapping the ratiometric emission ($R$) of ligand-binding two-state biosensors to their ligand concentration in their surrounding environment under equilibrium conditions. These maps follow the same form as those mapping $R$ to pH in two-state ratiometric pH biosensors. This enables us, in this note, to follow the scheme used in Supplementary Note 8, to model how the precision of $R$ measurements with a ligand-binding biosensor determines the range of ligand concentration values that is possible to measure with that biosensor at a specific inaccuracy level. We (9.1) estimate the values of the minimal set of parameters and constraints between parameters needed to predict the influence of relative errors in $R$ on ligand-binding biosensors with known $K_d$ and fluorescence spectra; and (9.2) determine the ligand-concentration inaccuracy we would expect to observe with each of those biosensors if we selected optimal excitation or emission filters for each biosensor and measured $R$ at the same precision as our $R$ measurements in the feeding muscles of live *C. elegans* with roGFP-R12.

**(9.1) Obtaining the values of the parameters that map $R$ to ligand concentration for ligand-binding biosensors**

<u>(9.1.1) Overall approach</u>

In Supplementary Note 7.2.2 we derived the function mapping a biosensor's fluorescence ratio ($R$) and the free ligand concentration in the environment surrounding the biosensor under equilibrium conditions.

$$[Ligand] = 10^{-pK_d + log_{10}(\delta_{\lambda_2}\frac{R_{Biosensor}-R}{R-R_{Biosensor-Ligand}})} \quad (1)$$

which can be rewritten as:

$$p[Ligand] = pK_d - log_{10}(\delta_{\lambda_2}\frac{R_{Biosensor}-R}{R-R_{Biosensor-Ligand}}) \quad (2)$$

This function has the same form as the function mapping $R$ and pH (see Supplementary Note 7.1.3) and therefore depends on equivalent biochemical and biophysical parameters. The biochemical parameter is the biosensor's dissociation constant $K_d$. This parameter is often known from published reports; however, in many cases the values reported do not represent true $K_d$ values as they are extracted from ligand-titrations of the biosensor's fluorescence ratio, which do not necessarily have a linear relationship with the fraction of ligand-bound biosensors (see Supplementary Note 2). Therefore, we corrected those effective $K_d$ values with knowledge of some of the biosensor's biophysical parameters, following the scheme described for pH biosensors in Supplementary Note 8.1.2.

Published reports often include the spectra only of biosensor species that are not fully ligand-bound and therefore underestimate the biosensor's dynamic range. We estimated biosensor dynamic range using published spectra obtained at two ligand concentrations and knowledge of the biosensor's $pK_d$ value, following the scheme described for pH biosensors in Supplementary Note 8.1.3.

We estimated the values of the minimal set of parameters and constraints between parameters needed to predict the influence of relative errors in $R$ on six ligand-binding biosensors with known $pK_d$ and fluorescence spectra (section 9.1.2). The value of $\delta_{\lambda_2}$ and the biosensor's overall dynamic range $DR$ (equal to the $R_{Biosensor}/R_{Biosensor-Ligand}$ ratio) was determined from the fluorescence spectra of ligand-bound and unbound biosensors. As discussed in Supplementary Note 5.1.3 for the map from $R$ to $E_{GSH}$, mapping the $R$ values obtained with a given instrument to $p[Ligand]$ requires knowledge of the values of both $R_{Biosensor-Ligand}$ and $R_{Biosensor}$ in that instrument; however, when all other parameters are known, knowledge of the $R_{Biosensor}/R_{Biosensor-Ligand}$ ratio is sufficient for modeling how relative errors in $R$ would influence the expected $p[Ligand]$ inaccuracy of different biosensors, under otherwise identical microscopy conditions.

(9.1.2) $pK_d$, $\delta_{\lambda_1}$, $\delta_{\lambda_2}$, $R_{Biosensor-Ligand}$ and $R_{Biosensor}$, and $DR$ parameter values for six two-state ratiometric ligand-binding biosensors

We attempted to obtain the biosensor's $pK_d$ value and the fluorescence excitation spectra of ligand bound and unbound biosensor species from published data for two-state ratiometric ligand-binding biosensors specific for histidine, $NAD^+$, NADH, and NADPH (Table S9-1). Of the nine ratiometric two-state ligand-binding biosensors, four had known $pK_d$ values (Table S9-1) and one had a known $pK_{effective}$ from which we estimated $pK_d$, following the scheme described in section 8.1.2 (Table S9-2). The fluorescence spectra of ligand-bound and unbound biosensor species have not been reported for four of the five iNAP biosensors (Table S9-1), including the iNAP1-

mCherry biosensor. For that biosensor, we assumed iNAP1 NADPH binding affinity and fluorescence emission at 535 nm was unaffected by fusion to mCherry.

For the five biosensors with available spectra and known or estimated $pK_d$ (FHisJ, NAD⁺ Biosensor, Frex, FrexH, iNAP1), spectra collected at different ligand concentrations were digitized with WebPlotDigitizer software [10], and the spectra of fully ligand-bound and fully unbound biosensor states were estimated following the scheme described in section 8.1.3. In all cases these spectra differed only slightly from the empirical measurements (Figure S9-1, first column).

We note that because these five biosensors consist of fusions of ligand-binding domains to circularly permuted fluorescent proteins (cpFPs), their spectral properties inherit the pH-dependence of their cpFPs domains. Typically, fluorescence in the first excitation band is pH-resistant, while fluorescence in the second excitation band is pH-sensitive (but with $pK_a$ insensitive to ligand binding). Parameter values were calculated from spectra acquired at a pH of 7.4 (Table S9-1). The iNAP1-mCherry biosensor allows ratiometric and pH-resistant measurement when using the pH-resistant 420 nm excitation band of iNAP1 [32], since mCherry red fluorescence is pH-resistant over a broad range of pH values because it has a $pK_a$ of less than 4.5 [33].

Parameter values for $\delta_{\lambda_1}$, $\delta_{\lambda_2}$, $R_{Biosensor-Ligand}$, $R_{Biosensor}$, and $DR$ were collected from the estimated fluorescence intensity spectra of fully ligand-bound and unbound biosensor states (Table S9-2). For each biosensor, we chose excitation- emission-wavelength pairs, $\lambda_1$ and $\lambda_2$, seeking to maximize $DR$ and to match the wavelengths at which biosensor fluorescence intensity peaked. We used biosensor fluorescence signals averaged over 20 nm intervals centered around $\lambda_1$ and $\lambda_2$ to calculate $\delta_{\lambda_1}$, $\delta_{\lambda_2}$, $R_{Biosensor-Ligand}$, $R_{Biosensor}$, and $DR$ values. The values of these parameters of each biosensor were determined as described in (9.1.2). These parameters varied significantly between biosensors (Table S9-2 and Figures S9-1 second column).

**(9.2) Predicted $p[Ligand]$ inaccuracy of six ratiometric two-state ligand-binding biosensors**

Using the $pK_d$, $\delta_{\lambda_2}$, $R_{Biosensor-Ligand}$, and $R_{Biosensor}$ parameters obtained in (9.1.2), we mapped $R$ to $p[Ligand]$ for six ratiometric two-state ligand-binding biosensors. We applied the SensorOverlord framework to determine the $p[Ligand]$ inaccuracy that we would expect to observe at each $p[Ligand]$ value with each of those biosensors, given our empirical distribution of relative errors in $R$ when measuring $E_{GSH}$ in the feeding muscles of live *C. elegans* expressing roGFP-R12 (Figure S9-1, third column).

#### Table S9-1: Two-state ratiometric ligand-binding biosensors

| Ligand | Biosensor | | $pK_d$ | $pK_{effective}$ | | Spectra? | |
|---|---|---|---|---|---|---|---|
| Histidine | FHisJ | [34] | 4.66 | | [34] | Yes | [34] |
| $NAD^+$ | $NAD^+$ Biosensor | [35] | 4.19 | | [35] | Yes | [35] |
| NADH | Frex | [36] | | 5.43 | [36] | Yes | [36] |
| NADH | FrexH | [36] | 7.40 | | [36] | Yes | [36] |
| NADPH | iNAP1 | [32] | 5.70 | | [32] | Yes | [32] |
| NADPH | iNAP2 | [32] | 5.22 | | [32] | No | |
| NADPH | iNAP3 | [32] | 4.60 | | [32] | No | |
| NADPH | iNAP4 | [32] | 3.92 | | [32] | No | |
| NADPH | iNAP1-mCherry | [32] | 5.70 * | | [32] | Inferred † | [32] |

* We assumed iNAP1 NADPH binding affinity was unaffected by fusion to mCherry.

† We assumed iNAP1 flurescence emission at 535 nm was unaffected by fusion to mCherry.

### Table S9-2: Biosensor parameter values used for mapping $R$ to ligand concentration

| Biosensor | Imaging modality | $\lambda_1 \pm 10\,nm$ | $\lambda_2 \pm 10\,nm$ | $pK_d$ | $DR$ | $\delta_{\lambda_1}$ | $\delta_{\lambda_2}$ | $R_{Bios-Ligand}$ | $R_{Biosensor}$ |
|---|---|---|---|---|---|---|---|---|---|
| FHisJ | dual excitation | 415 | 485 | 4.66 | 6.05 | 2.06 | 0.34 | 0.32 | 1.93 |
| $NAD^+$ Bio-sensor | dual excitation | 500 | 425 | 4.19 | 1.84 | 1.86 | 1.01 | 2.04 | 3.76 |
| Frex | dual excitation | 410 | 500 | 5.45 | 4.84 | 0.93 | 0.19 | 0.15 | 0.73 |
| FrexH | dual excitation | 500 | 410 | 7.40 | 1.65 | 2.00 | 1.22 | 0.92 | 1.51 |
| iNAP1 | dual excitation | 500 | 420 | 5.70 | 7.45 | 2.13 | 0.29 | 0.26 | 1.93 |
| iNAP1-mCherry | dual excitation / dual emission | 560 / 630 | 420 / 535 | 5.70 * | 3.5 † | 1 † | 0.29 † | 1 † | 0.29 † |

* We assumed iNAP1 NADPH binding affinity was unaffected by fusion to mCherry.

† We assumed iNAP1 flurescence emission at 535 nm was unaffected by fusion to mCherry.

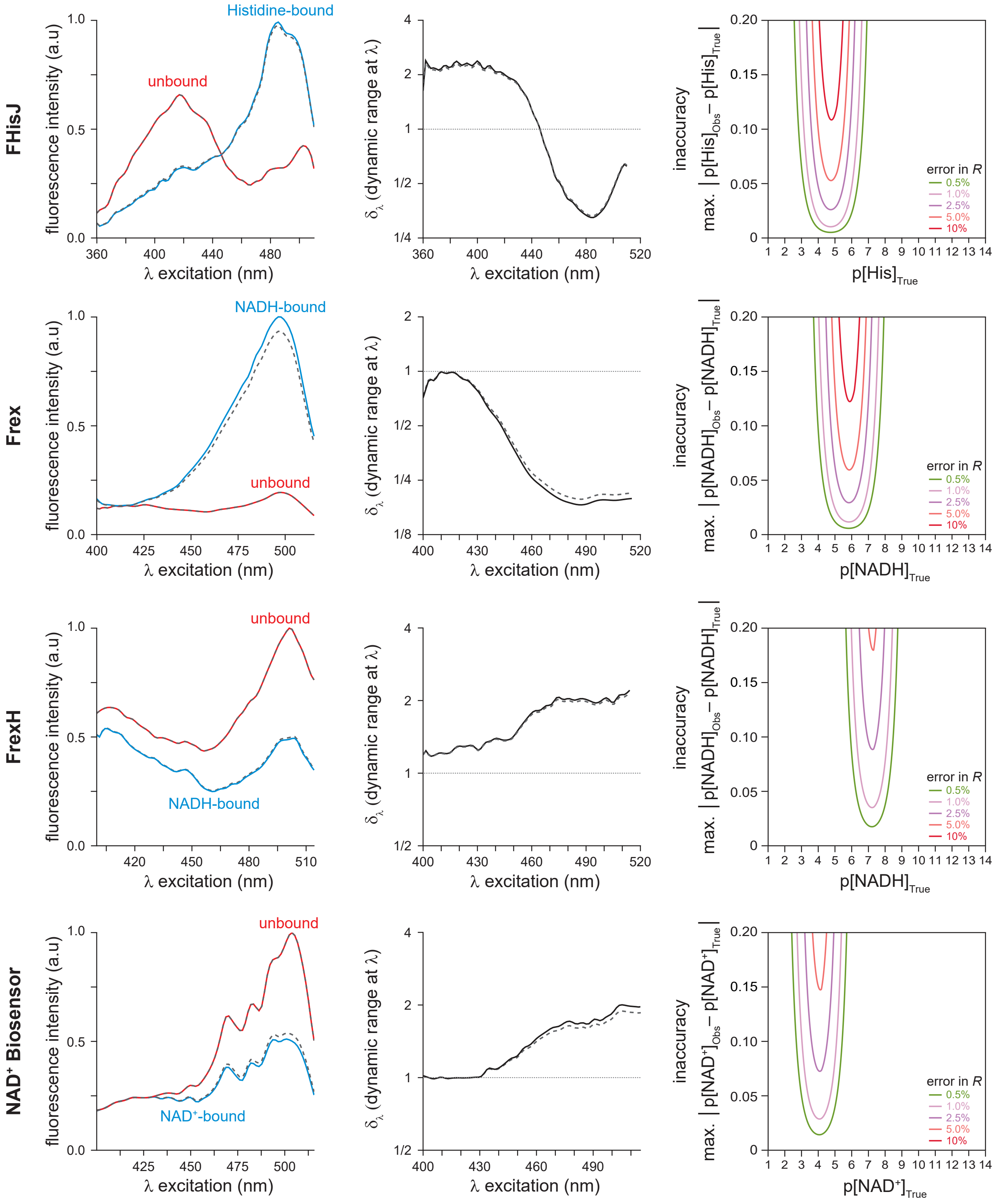


**Figure S9-1: Predicted $p[Ligand]$ inaccuracy of ratiometric two-state ligand-binding biosensors**

In this analysis of five dual-excitation ligand-binding biosensors, and one dual-excitation dual-emission ligand-binding biosensor, we show in each row a plot of the biosensor's predicted emis-

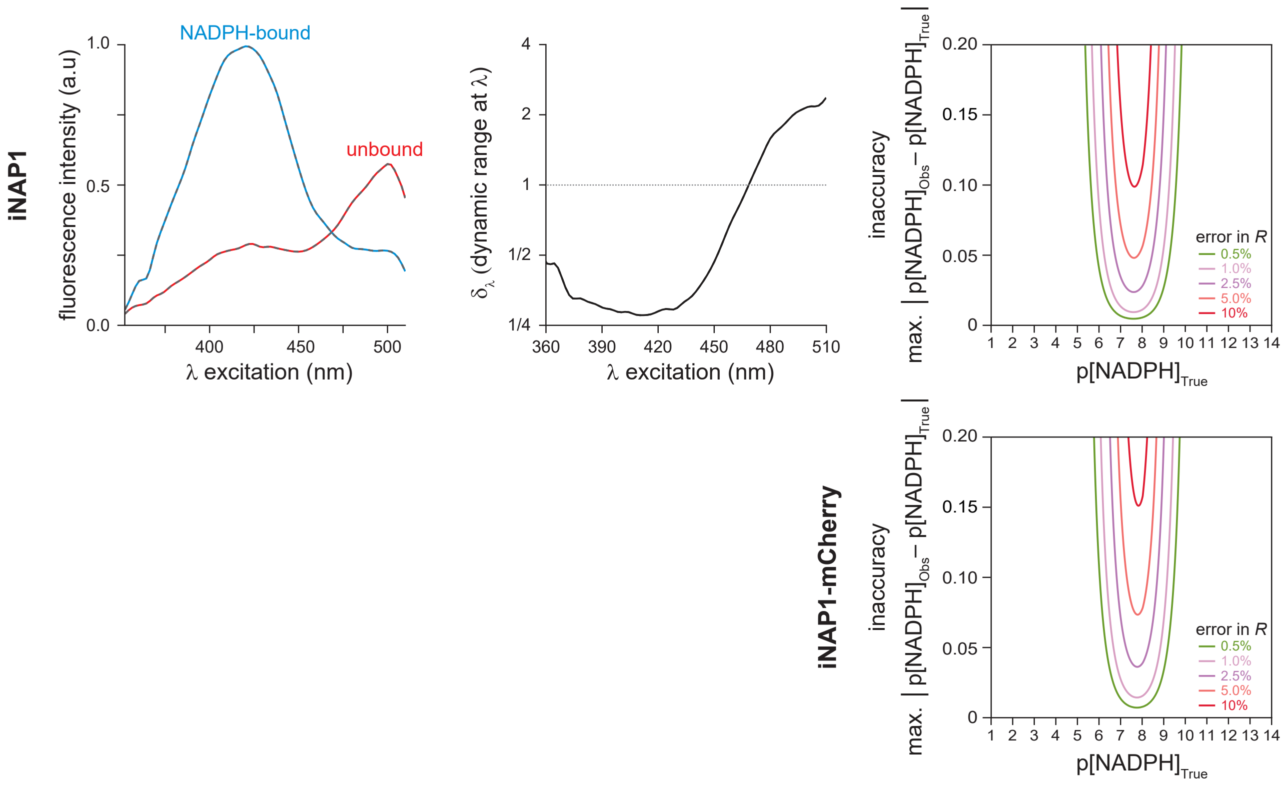


sion when excited at a wavelength $\lambda$ in the fully ligand-bound and unbound biosensor states (solid lines, column 1) and the observed emissions of mostly ligand-bound and mostly unbound biosensor mixtures (dotted lines, column 1); a plot of the biosensor's dynamic range $\delta_\lambda$, the predicted ratio of the emissions of the fully ligand-bound and unbound biosensor states (solid line, column 2) and the apparent $\delta_\lambda$ derived from the ratio of the observed emissions of mostly ligand-bound and mostly unbound biosensor mixtures (dotted line, column 2); and a plot of the predicted $p[Ligand]$ measurement inaccuracy with that biosensor at each wavelength $\lambda$ for different relative errors in $R$ values (column 3).
